# A hierarchy of time constants and reliable signal propagation in the marmoset cerebral cortex

**DOI:** 10.1101/2025.05.18.654729

**Authors:** Guanchun Li, Songting Li, Xiao-Jing Wang

## Abstract

A hierarchy of timescales in the cerebral cortex is functionally desirable for rapid information processing in sensory areas and slow time integration in association areas. Here, through an analysis of electrocorticography (ECoG) data, we identified a timescale hierarchy in the neocortex of marmoset, a primate species commonly used in neuroscience. Constrained by the anatomical and electrophysiological data, we developed a multi-regional model of the marmoset neocortex that captures the observed timescale hierarchy phenomenon. Furthermore, we used the model to investigate how the neocortex reconciles information integration on distinct timescales in local areas with reliable signal propagation globally across regions. We found that a near-criticality state is optimal for both localized signal integration within areas and reliable signal propagation across areas in the multi-regional neocortex. Our model also mechanically explains recent experimental observations that the structural and functional connectivities are less correlated in association areas than in sensory areas.

## 1 Introduction

The cerebral cortex, together with peripheral sensory organs (e.g., retina, cochlea, spinal cord) and subcortical relays (e.g., thalamus), engages in information processing of external inputs, local integration of signals within regions, and the transmission of these signals along functionally specific pathways. A prominent and noteworthy characteristic of the neocortex is what is referred to as “timescale hierarchy”. In essence, different cortical areas exhibit varying timescales in their response to inputs, allowing for the hierarchical processing of temporal information. Primary sensory areas manifest shorter timescales, typically measured in tens of milliseconds. The short timescales enable these areas to respond rapidly and effectively to incoming stimuli. In contrast, higher-level association areas display longer timescales, often extending to hundreds of milliseconds or longer. These extended timescales support the integration of information over a relatively protracted time window. The phenomenon of timescale hierarchy emerges from a theoretical model of the multi-regional macaque neocortex [1], and is supported by experimental studies conducted on multiple species, including mice [2, 3], macaque monkeys [4, 5], and humans [6–11]. It remains to be actively investigated on the universality of this phenomenon across different species and the corresponding large-scale circuit mechanisms.

Timescale hierarchy is a phenomenon that spans across multiple cortical areas, strongly suggesting that its underlying mechanism is intricately linked to the broadscale structural attributes of the neocortex. On an intuitive level, the timescale of neuronal population activity is intricately tied to the strength of recurrent excitation within that population. In simpler terms, stronger recurrent excitation tends to result in slower timescales of activity. Consequently, the concept of timescale hierarchy points to the existence of a macroscopic gradient in recurrent excitation along the axis of cortical area hierarchy. This notion finds compelling support in experimental findings, particularly in studies that examine the number of dendritic spines on pyramidal neurons within the primate neocortex. These dendritic spines serve as a direct reflection of the synaptic excitation strength per neuron and notably display an increasing gradient along the cortical hierarchy [12, 13]. Furthermore, beyond this spatial gradient in spine numbers, the neocortex also reveals macroscopic gradients in other biological properties, including neuron density [14] and synaptic receptor distribution [15, 16]. These gradients may indeed serve as fundamental principles governing the organization of the cerebral cortex on a large scale, as discussed in a comprehensive review [17].

In addition to the inherent characteristics of individual cortical areas, timescale hierarchy may also be attributed to long-range connections among these areas. Recent years have witnessed significant advancements in the quantitative measurement of inter-areal connectivity in various species, including mouse [18–21], macaque [22–24], and marmoset [25, 26]. It represents a departure from earlier studies that only provided a coarse assessment of primate connectomes by categorizing connections between area pairs as absent, weak, or strong [27, 28]. In contrast, the contemporary connectome offers a more precise and quantitative characterization. The strength of inter-areal connections is characterized using a continuous metric known as the fraction of labeled neurons (FLNs). FLNs measure the relative strength of projections from a particular source area to a specific target area in relation to all source areas projecting to that target area. Such a detailed connectome framework enables the quantitative exploration of the organizational principles governing the cerebral cortex. For instance, one notable observation is that FLNs between two areas exhibit an exponential decay with increasing distance, a phenomenon known as the exponential distance rule (EDR) [29, 30].

By incorporating the macroscopic gradient of excitation and a quantitative weighted and directed connectome of macaque monkey, a comprehensive multi-regional model of 29 areas was developed to qualitatively generate the timescale hierarchy phenomenon [1]. Notably, when the macroscopic gradient of excitation was removed or the connectivity was shuffled, the significance of the timescale hierarchy diminished, or in some instances, completely vanished [1]. To elucidate the underlying principles governing this phenomenon, a rigorous mathematical analysis was conducted to identify the importance of macroscopic gradient of excitation and detailed excitation-inhibition (E-I) balance. The balance condition stipulated that the inter-areal excitatory inputs originating from each source area must balance with the local inhibitory inputs from the target area [31]. As a result, cortical areas exhibited relatively weak effective interactions, leading to the emergence of timescale hierarchy in a densely connected network.

While this detailed balance condition appeared to enhance signal gating, i.e., irrelevant signals are not able to propagate in the detailed balance state, as demonstrated in a computational study [32], it also introduced a challenge: the model struggled to achieve reliable signal propagation, another crucial dynamic property of the large-scale neural circuits. On the other hand, signals experienced significantly less attenuation when they operated within an alternative dynamical regime known as “global balanced amplification” [33]. However, in this regime, the timescale hierarchy phenomenon ceased to exist. The conundrum lies in reconciling both timescale hierarchy and reliable signal propagation within the cerebral cortex. Furthermore, despite the model being informed by anatomical data, it remains unclear to which extent the model accurately captures the timescales of individual areas and the precise properties of signal propagation in a quantitative manner. A challenge stems from the absence of neuronal activity data recorded across multiple areas in the macaque neocortex.

In this study, starting from analyzing the multi-regional electrocorticography (ECoG) recording data of another primate, the marmoset, we investigated two critical aspects within the marmoset neocortex: the timescale hierarchy phenomenon and the property of signal propagation. It is notable that there has been limited prior research reporting the existence of timescale hierarchy in the marmoset neocortex. Consequently, it becomes imperative to explore whether this phenomenon also manifests in the marmoset, thereby testing the hypothesis that timescale hierarchy may serve as a universal principle governing cortical dynamics across different species. In addition, the availability of comprehensive anatomical data [25, 26] along with activity data [34] collected from multiple cortical areas within the marmoset neocortex, provides us with an opportunity to develop a biologically realistic large-scale model that encompasses multiple cortical areas. Importantly, and in contrast to previous research, the predictions made by our model regarding the timescales of individual cortical areas and the patterns of signal propagation across multiple areas can be directly validated using experimental activity data. This anatomically and dynamically constrained model empowers us to delve deeply into the role played by the intra-areal heterogeneity (i.e., macroscopic gradient of excitation) and inter-areal connectivity in shaping local signal integration and global signal propagation dynamics within the marmoset neocortex, in particular, leading to a near-critical state of the neocortex. In addition, the model mechanically explains recent experimental observations that the structural connectivity and functional connectivity are less similar in association areas than in sensory areas.

## 2 Results

### 2.1 Experimental evidence of timescale hierarchy in the marmoset neocortex

We initiated our investigation with the analysis of an ECoG dataset recorded from an anesthetized marmoset in a previous study [34]. This dataset includes signal measurement from 96 electrodes, broadly placed on the left hemisphere of the marmoset’s brain during a state of passive listening (Figure 1a). To circumvent any potential bias from the auditory stimulus, we selected data recorded prior to any auditory stimulus exposure, which was presumably close to the resting state (for further details, see Sec. Methods).

**Fig. 1.**
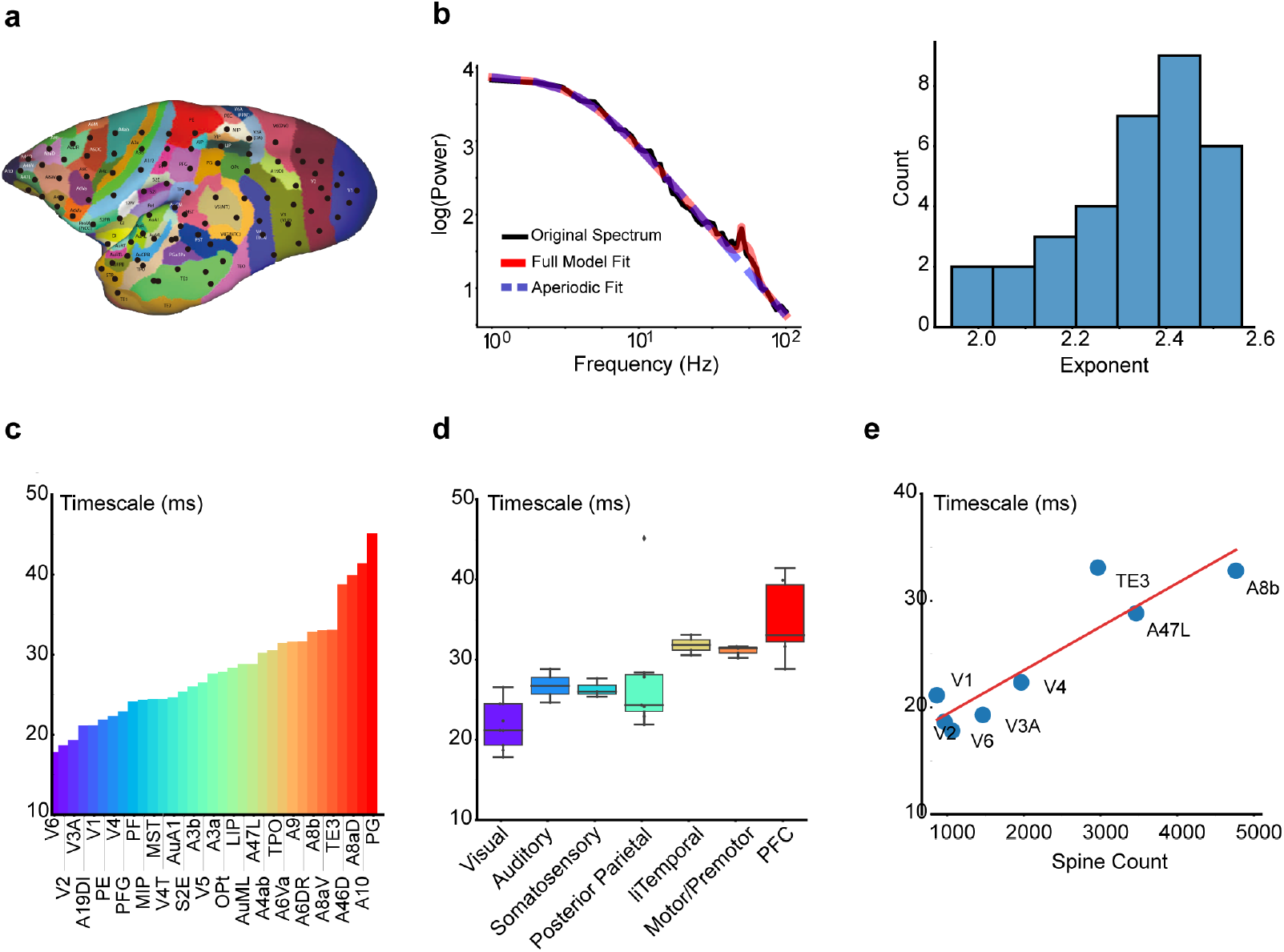
Experimental observation of timescale hierarchy in the marmoset neocortex. (a) The spatial distribution of electrode locations marked by black circles. (b) Methodology for timescale measurement. Left panel: Exemplification of the Power Spectral Density (PSD) fitting process. Right panel: Distribution of the exponent values derived from the Lorentz function fit of the power spectrum. (c) Timescale distribution across the marmoset neocortex. Lower-level sensory areas such as V1, V2, and AuA1 exhibited shorter timescales, in contrast to higher-level areas like A8aD, A10, and PG that exhibited longer timescales. (d) Boxplot of timescale statistics for each areal category. Box plots show the median and IQR, with whiskers extending to the minimum and maximum values excluding outliers. (e) Positive correlation between timescale and spine count of pyramidal neurons for each area, revealing the structural-dynamical relationship (*r* = 0.91, *p* = 1.91 *×*10^−3^). The timescales are fitted using the experiment data from an anesthetized marmoset [34].

We adapted a spectral approach to estimate the timescales of resting-state neural activity from the ECoG dataset [11]. This method focuses on the Power Spectral Density (PSD) of the data, enabling reliable estimations of the time constant of neural activity from relatively short time series. In brief, we calculated the PSD of the activity data using Welch’s method, followed by a Lorentz function fitting to determine the ‘knee frequency’ and the corresponding decay time constant (see details in Sec. Methods). Previous studies anticipated an exponent of 2 for the Lorentz function [35], which has also been observed in our study showing that the exponent clustered around 2 with minor variance in Figure 1b.

By applying the spectral method, a prominent timescale gradient across 33 cortical areas was revealed. As illustrated in Figure 1c, sensory areas including V1, V2, and AuA1 exhibited smaller time constants, while association areas including A8aD, A10, and PG exhibited longer timescales. This trend was further substantiated by computing the average timescale for each category of areas, presenting a significant increase of timescales from the low-level sensory areas to the high-level association areas, as shown in Figure 1d.

Intriguingly, our data analysis unveiled a striking consistency between the timescale hierarchy and the structural hierarchy assessed by the spine number of pyramidal neurons in each area [1, 12]. As demonstrated in Figure 1e, the timescale correlates strongly with the spine count across cortical areas. Furthermore, by analyzing the first 5 seconds of awake resting-state recordings [36], we observed a clear intrinsic timescale hierarchy across cortical areas closely matching the anesthetized state (Figure S9a-b), with a strong correlation confirming consistency in the relative ordering of timescales (Figure S9c).

Those findings propelled us to construct a multi-regional model of the marmoset neocortex that links the structural and dynamical property of the neocortex. With such a model, we were able to investigate how network structure shapes signal integration reflected by the neural time constant within each area and signal propagation across areas in the cerebral cortex – in particular, the role of the macroscopic gradient of spine count and the inter-areal connectivity.

### 2.2 A multi-regional model of the marmoset neocortex

We next developed a multi-regional model of the marmoset neocortex. The form of the model was adapted from Ref. [1] that described the macaque neocortex, considering the anatomical resemblance between the macaque and marmoset cortices. The model encapsulates 55 distinct sensory and association areas of the marmoset neocortex, including the 33 areas recorded in the previous experiment [34]. Each area in our model encompasses an excitatory and inhibitory neural population, both of which are locally interconnected with their activity described by the population firing rates. A detailed schematic of this multi-regional model is presented in Figure 2a-b.

**Fig. 2.**
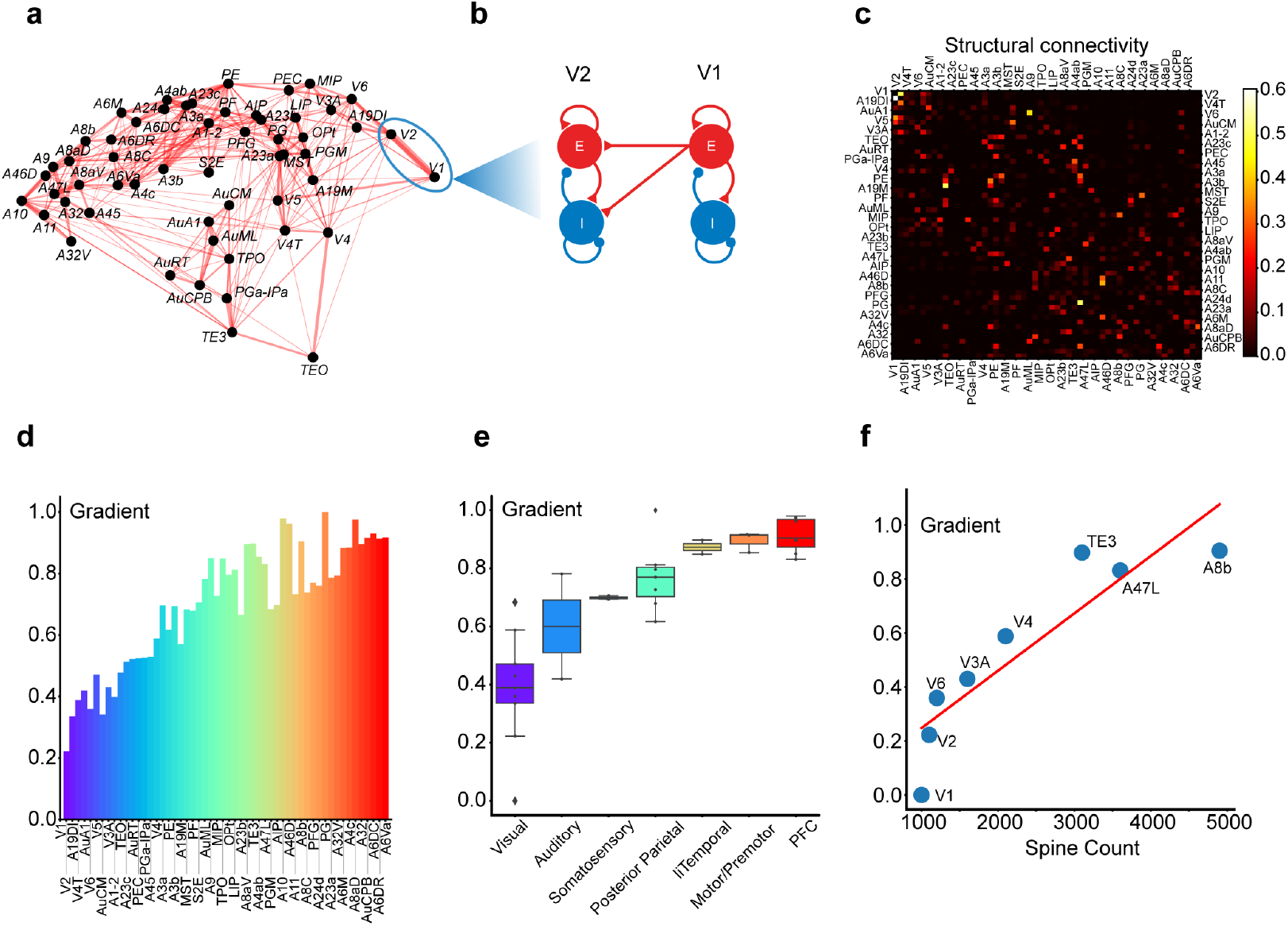
Properties of the multi-regional model of the marmoset neocortex. (a) Graph representation of the marmoset cortical areas, with line width representing connection strength (only the strongest 25% of connections are displayed). (b) Schematic depiction of the model area representing each cortical area, comprising excitatory and inhibitory neural populations with the directed interaction from Area V1 to Area V2 shown. (c) Matrix of connection strengths across all cortical areas. The directed connection magnitude between any two areas is based on the FLN data derived from [25, 26]. Only inter-areal (long-range) projections are shown for clarity. (d) Area-wise composite gradient of excitation in the model. (e) Boxplot of the composite gradient of excitation grouped by the category of areas, signifying a significant progression from visual areas to the prefrontal cortex. Box plots show the median and IQR, with whiskers extending to the minimum and maximum values excluding outliers. (f) Strong correlation between the model’s composite gradient of excitation and the experimentally measured spine number of 8 areas (*r* = 0.89, *p* = 2.6 *×*10^−3^). Layer-3 pyramidal-cell spine counts were taken from the cross-area compilation in Supplementary Table 4 of [26]; sampling details are given in the cited source articles.

The network connectivity of the model was determined by the up-to-date anatomical measurement using retrograde tracing [25, 26]. The magnitude of the directed connection between any two areas was derived from FLNs determined through tracer injection experiments. The FLNs quantify the fraction of neurons projecting to a target area across all source areas, which can be viewed as a proxy of connectivity strength between two areas in our model. A heatmap representation of the FLN matrix is shown in Figure 2c. The dynamical interactions within each area are described following a Wilson-Cowan type model (see further details in Sec. Methods).

In addition, the model accentuates the influence of the excitation gradient in scaling connection strengths between neural populations. Following prior macaque neocortex modeling [1], we integrated a composite gradient of excitation level across areas in the model, which mainly reflects the areal specific spine numbers of pyramidal neurons based on a previous anatomical study [26]. In our model, this composite gradient index has been additionally constrained to align with the experimentally measured timescales deduced from the ECoG data, as detailed in the preceding section. A detailed methodology for estimating the composite gradient is described in Sec. Methods. The composite gradient’s barplot is shown in Figure 2d, alongside the average composite gradient for each category of cortical areas in Figure 2e, revealing a statistically significant upward trend from visual areas to the prefrontal cortex. Furthermore, Figure 2f illustrates the correlation between the composite gradient and the spine number of pyramidal neurons for all areas [26], demonstrating that the composite gradient index was predominantly determined by the spine number of pyramidal neurons, thus as a proxy of excitation level for neurons in each area.

While subcortical structures (e.g., thalamus, brainstem, basal ganglia, cerebellum) and peripheral sensory pathways (e.g., retina, cochlea, spinal cord) exert influences on cerebral cortical dynamics, the present model is restricted to cerebral cortical networks due to the absence of a marmoset-wide, high-resolution subcortical and peripheral connectome. It is worth noting, however, that this work focuses on timescales of spontaneous neural activity during the resting state, when there is no experimentally designed sensory stimulation to peripheral structures and animals are not engaged in a particular task. As future anatomical data allow the integration of subcortical and peripheral circuits, these additional components are expected to improve our model predictions of timescales and signal propagations, with the specific patterns likely influenced by interactions involving subcortical and peripheral structures.

### 2.3 Emergence of timescale hierarchy in the marmoset network model

We next investigated the capability of our model in replicating the experimentally observed timescales for each area in the marmoset neocortex. In simulating the model’s resting state, we applied white-noise stimuli across all brain areas, to emulate the resting state scenario in experiment. The neural activities in various areas, illustrated in Figure 3a, reveal that lower-level areas resonate at higher frequencies, akin to the input white noise. In contrast, higher-level areas exhibit smoother variations, reflecting their differing operational timescales. Further inspection of the auto-correlation function (ACF) of activity within each area, as shown in Figure 3b, demonstrates an evident exponential decay, though with significant variability in the decay time constants across areas, thus reinforcing the existence of different operational timescales in each area.

**Fig. 3.**
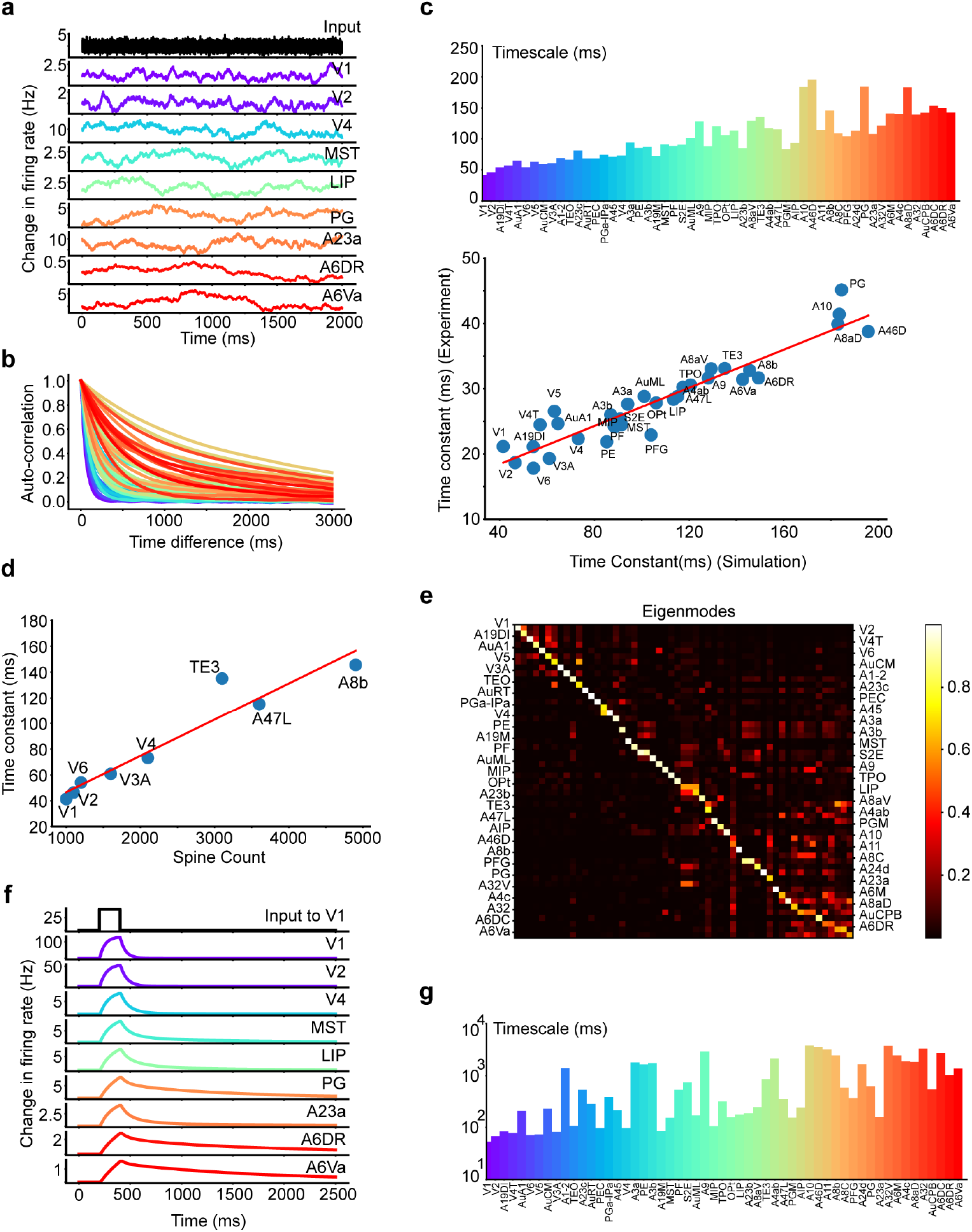
Investigation of timescale gradient and localization in the model. (a) Simulated resting state activity, with white-noise input to all brain areas. Lower-level areas resonate at higher frequencies, whereas higher-level areas show more gradual fluctuations. (b) The autocorrelation function of each area’s activity in the resting state. (c) Top panel: The timescale derived from the Power Spectral Density (PSD) of simulated resting state activity, forming a gradient spanning from 50ms to 250ms. Bottom panel: Strong correlation of timescale between model and experiment (*r* = 0.94, *p* = 6.57 *×*10^−16^). (d) Significant correlation between the simulated timescale and spine number (*r* = 0.95, *p* =2.18*×* 10^−4^). (e) Visualization of the dynamical system eigenmodes. The timescale localization for each area is portrayed by the sparseness of the eigenmodes. (f) The activity of selected areas following a stimulus to V1 in the model. Decay time constant increases along the signal propagation pathway. (g) The observed timescale gradient following a stimulus to V1, as extracted by fitting to the exponential function.

We then examined the timescale gradient more closely by estimating timescales from the PSD of the simulated neural activity, following the same approach used for the experimental data processing. As displayed in Figure 3c, the timescale for each area forms a gradient ranging from 50ms to 250ms. This gradient rises from sensory to high-level areas, echoing the experimentally observed pattern. Next, we quantitatively compared the experimentally measured and model predicted timescales area by area. Figure 3c shows a strong correlation between them, with a scaling factor around 5. This scaling is consistent with the findings in Ref. [11] where the timescale of ECoG was compared with the timescale of neuronal firing activity. These outcomes suggest our model well captures the experimental observations, thus validating the model’s effectiveness. Moreover, the timescale gradient and its close consistency with experimental data remain robust even when biologically realistic axonal conduction delays are incorporated into the model (Figure S8). Finally, Figure 3d demonstrates that the consistency between the time constant and the spine number across areas is also preserved in our model.

Furthermore, we substantiated the property of timescale gradient by examining the eigenmodes in the modeled system. For each eigenmode (a column of the matrix in Figure 3e), each entry implies that the neuronal activity in the corresponding area will possess a timescale determined by the eigenvalue of that eigenmode. Thus, a sparse eigenmode indicates that only a few areas exhibit a particular timescale determined by the corresponding eigenvalue, supporting the concept of “timescale localization”. Figure 3e illustrates the phenomenon of timescale localization as indicated by the eigenmode matrix’s resemblance to a block diagonal matrix. Such an observation is instrumental as it indicates the capacity for lower-level sensory areas to respond quickly to incoming stimuli and discard older inputs. Conversely, higher-level planning and decision-making areas retain information over extended periods, aiding in integrative processes such as working memory.

We further studied the timescale gradient in the presence of stimulus in the model. By recording the post-stimulus response for all areas after activating the V1 area, Figure 3f demonstrates that activity in high-level areas (PG, A6DR, A6VA, etc.) persisted longer than in lower-level visual areas (V1, V2, V4, etc.), reflecting a functional facet of the timescale gradient. A further quantitative investigation of this gradient by an exponential fit to the activity data (details in Sec. Methods) was depicted in Figure 3g. Intriguingly, the auditory areas, typically considered lower-level but not directly involved in visual stimulus transmission, showed a longer timescale. This observation can potentially be attributed to the absence of direct connections between visual and auditory cortices. Hence, when the visual cortex is stimulated, the auditory cortex may be indirectly activated through high-level cortical areas operating over longer timescales. As a result, the auditory cortex, receiving inputs over extended periods, also exhibits these longer timescales, as depicted in Figure 3g.

Figure S1a presents a direct comparison of timescales during rest and post-V1 stimulus, revealing a marked similarity between the two timescales derived under different brain states. Notably, higher-level areas exhibit longer timescales than lower-level ones, with exceptions like the auditory cortex discussed previously. We attribute these differences to the external stimuli on specific brain areas. Further mathematical analysis and numerical simulations (Supporting Information and Figure S1b-e) clarify that the post-stimulus timescale of area activity is influenced by the system eigenmodes’ intrinsic timescales and the stimulus-induced multi-regional activity pattern. Importantly, the external stimulus does not alter the network’s intrinsic timescale or eigenmode pattern but modifies the contribution of each intrinsic timescale (and the corresponding eigenmode) to the activity of cortical regions.

### 2.4 Essential conditions on timescale localization

We next studied the mechanism underlying the property of timescale localization. A previous theoretical study of the macaque neocortex [31] has proposed essential conditions for achieving timescale localization: the macroscopic gradient of excitation across areas and the balance of excitatory and inhibitory inputs received by each area. The macroscopic gradient refers to the gradual varying excitation strength across different cortical areas indicated by spine number of pyramidal neurons in each area, while ‘E-I balance’ alludes to the delicate current cancellation between global longrange excitatory inputs and local inhibitory synaptic inputs to the excitatory neural population in each area.

To examine these two conditions in the marmoset cortical network, we first introduced two metrics for eigenmodes – the Inverse Participation Ratio (IPR) and the *θ* index to quantify the timescale localization and its spatial localization, respectively (see both definitions in Sec. Methods, *θ* was initially proposed in Luis Carlos Garcia and Xiao-Jing Wang, see [37]). A higher IPR value (closer to 1) indicates a particular timescale localized in fewer number of areas, and a higher *θ* index (closer to 1) describes a timescale localized in physically more proximal areas. For the model detailed previously, the scatterplot of IPR and *θ* for each eigenmode of the model system is shown in Figure 4a, while the timescales of each eigenmode is indicated by the color code. Examples of eigenmodes with different IPR and *θ* are presented in Figure S2. As shown in Figure 4a (left panel), both high IPR and *θ* are observed for most eigenmodes, demonstrating that the timescale localization is present in the model, and each timescale is spatially localized in physically neighboring areas.

**Fig. 4.**
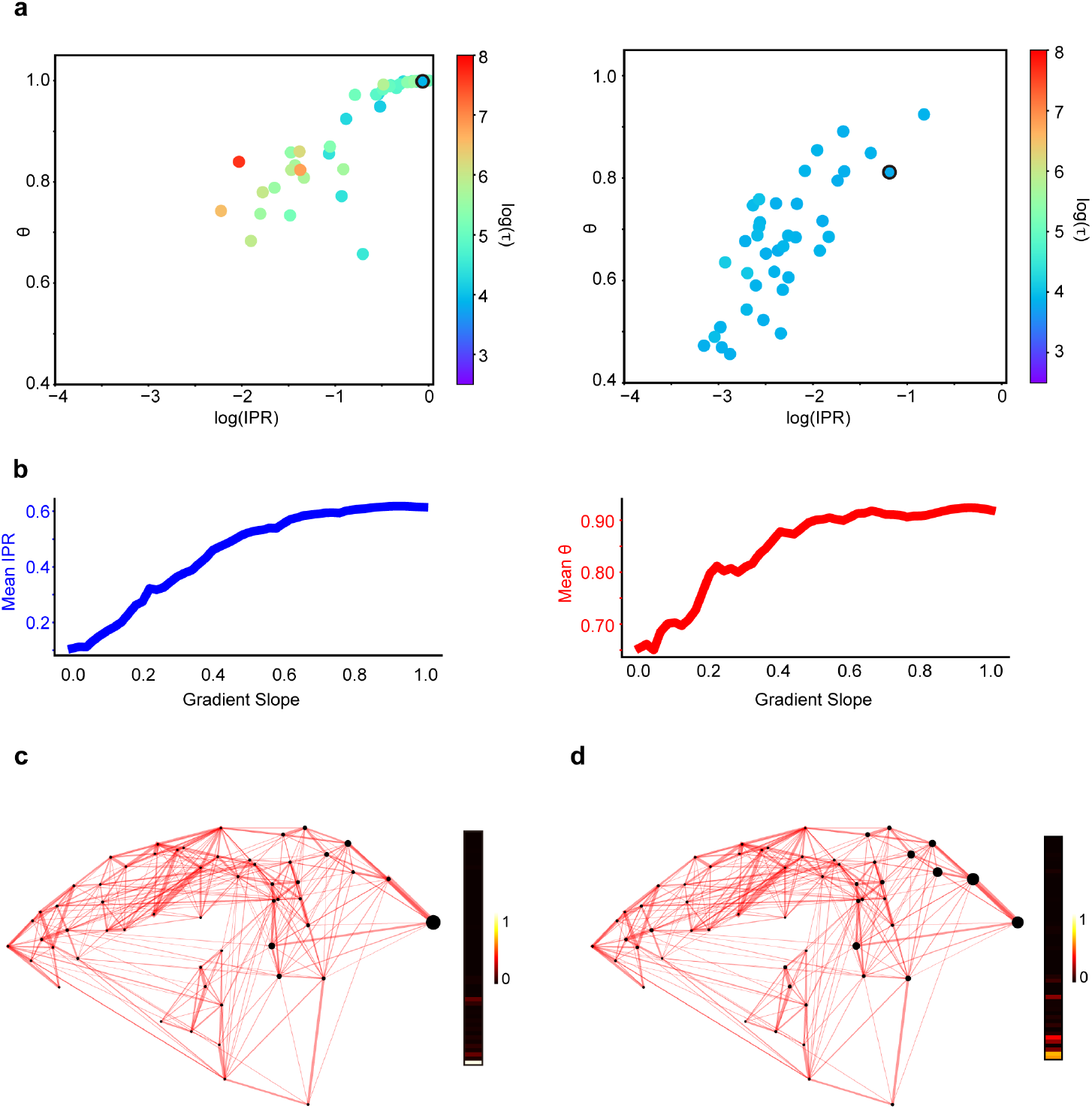
Effect of macroscopic gradient of excitation on timescale localization. (a) Scatterplot of IPR and *θ* for each eigenmode of the modeled dynamic system. Left panel: plot under the control condition. Right panel: plot under the condition that the composite gradient of excitation is absent. The colorcoded timescale was after taken the logarithm using the natural base e. (b) Change of IPR (left panel) and *θ* (right panel) as the gradient slope increases. (c) Visualization of a representative eigenmode (the fastest eigenmode) in the control condition, with values of the eigenmode plotted aside. The size of each black dot codes the magnitude of the corresponding element in an eigenmode. (d) Similar to (c), under the condition that the composite gradient of excitation is absent. The eigenmodes correspond to the black-circled dot in (a).

Subsequently, we examined the conditions for timescale localization in the marmoset neocortex. By removing the macroscopic composite gradients (setting the gradients *h*_*i*_ to zero for all areas in the model), we observed a marked degradation of timescale localization and spatial localization, as evidenced by significantly smaller IPR and *θ* (Figure 4a, right panel). Notably, the range of eigenmode timescales becomes narrower and more concentrated (predominantly around 50 ms), underscoring the influence of the composite excitation gradient in extending timescales. Further exploration involved adjusting the gradient’s slope (multiplying the gradient *h*_*i*_ with a factor *γ* for all areas in the model) and observing the resulting changes of timescale localization. We found a progressive increase in IPR from 0.1 to 0.65 and *θ* from 0.65 to 0.95 as the gradient slope *γ* varies from 0 to 1 (Figure 4b), supporting the pivotal role of the macroscopic gradient of excitation in timescale localization and its spatial distribution. The effect of the gradient of excitation becomes even more apparent when comparing the eigenmodes of the network with and without the gradient. Figures 4c and 4d showcase the fastest eigenmode for both scenarios (edged in black in Figure 4a). While both eigenmodes predominantly involve low-level visual areas with V1 activity dominating the eigenmode, the eigenmode in the control condition is markedly sparser, highlighting the dominant role of V1 activity. This comparison demonstrates the composite gradient’s contribution to timescale localization, even within the low-level sensory cortex.

In a parallel model simulation, we disrupted the E-I balance in our model by increasing the strength of local connections from inhibitory to excitatory neurons (*w*_*EI*_). This modification leads to the imbalance between the global long-range excitatory inputs and the local inhibitory synaptic inputs. Consequently, it resulted in smaller IPR and *θ* that reflect poor timescale localization and spatial localization (Figure 5a), similar to the case of gradient removal. With the E-I balance disturbed, the eigenmode characterized by the shortest timescale no longer involved the primary visual cortex (Figure 5b). This shift underscores the critical function of E-I balance, not merely as an essential condition for timescale localization but also as a determinant in the gradient of timescales from low-level sensory to high-level brain areas.

**Fig. 5.**
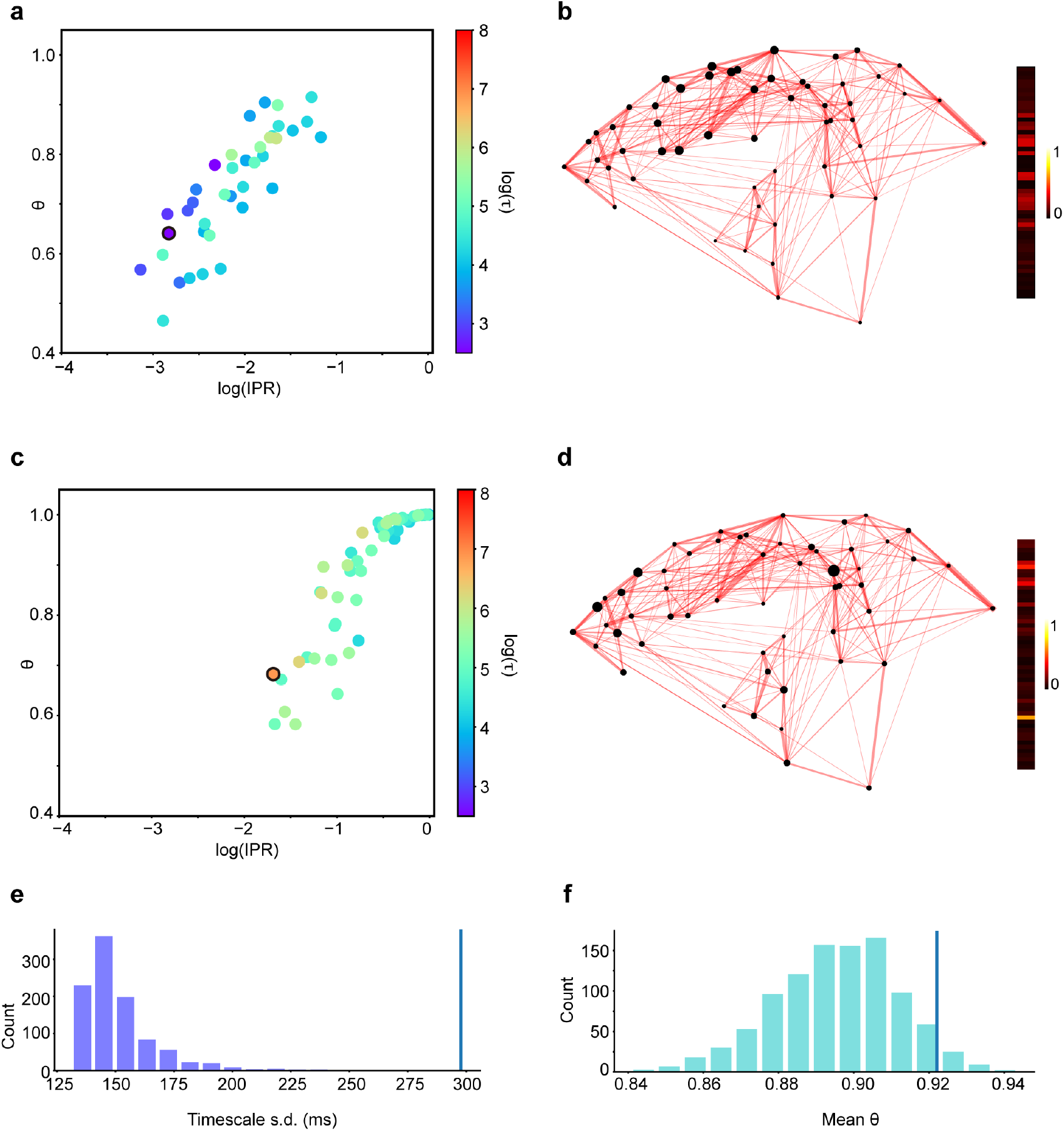
Essential conditions of timescale localization. (a) Scatterplot of IPR and *θ* for each eigenmode of the modeled dynamical system similar to Figure 4a, under the condition that the E-I balance is disrupted by increasing *w*_*EI*_ by 10%. (b) Visualization of the fastest eigenmode (black-circled dot in (a)), alongside the values of eigenmode, under the condition that the E-I balance is disrupted by increasing *w*_*EI*_ by 10%. (c) Similar to (a), but the FLN matrix is shuffled. (d) Similar to (b) but showing the slowest eigenmode (black-circled dot in (c)) with the shuffled FLN matrix. (e) Impact of shuffled FLNs on timescale range. The histogram of the timescale range quantified by the standard deviation of timescales after randomly shuffling FLNs. (f) Impact of shuffled FLNs on spatial localization. The histogram of the degree of spatial localization quantified by the *θ* metric with randomly shuffled FLN. Both distributions ((e) and (f)) are significantly smaller than the values observed in the control condition, as indicated by the vertical lines.

In our continued exploration, we probed the influence of long-range connections on timescale localization. When connectivity strengths (FLNs) were randomly shuffled, as illustrated in Figure 5c, the cortical network preserved high IPR values similar to the control condition (Figure S3), indicating robust timescale localization. However, *θ* index reduced significantly, suggesting a decrease in spatial localization of timescales. This effect is also evident using the example of the slowest eigenmode in both the shuffled FLN network and the control condition, as illustrated in Figure 5d and Supplementary Figure S2e, respectively. Notably, while both eigenmodes encompass a variety of brain regions, including notably the prefrontal cortices, the eigenmode under the shuffled FLN condition exhibits a broader distribution, extending to primary visual and auditory cortices. In contrast, the eigenmode in the control condition demonstrates greater spatial localization, predominantly engaging high-level brain areas.

Besides the reduced spatial localization, the range of timescales was also narrowed compared to the control condition. These observations were further substantiated by the results from 1000 repetitions of FLN shuffling. Disrupting these long-range connections led to a noticeable contraction in the timescale range and diminished spatial localization, as evidenced in Figure 5e, 5f. These results underscored the significant role of long-range connections in timescale localization, i.e., nearby areas process signals with similar timescales reflected by high *θ* value, while areas with some distance possess a variety of timescales enabling hierarchical signal processing. It suggests that the organization of the inter-areal connections is not random but is rather essential for brain functionalities.

### 2.5 Impact of near-critical state on structural and functional connectivity

As the macroscopic gradient of excitation largely influences timescales across areas, it is expected to influence the functional connectivity (FC) that relies on the temporally co-activation of pairwise areas. On the other hand, the macroscopic gradient also adjusts the effect of structural connectivity (SC) as the FLN will be scaled by the composite gradient as a proxy of the spine number of neurons in each area. Therefore, in this section, we direct our attention to how the macroscopic composite gradient shapes the correlation between SC and FC. We initiated this investigation by removing the gradient from our model and computing the resultant FC. Under these conditions, the simulated FC from our model significantly aligns with the SC. This increased correlation, contrasted with the dissimilar case in the presence of gradient of excitation (visualized in Figure 6a), suggests that the composite gradient can contribute to high-order connections within FC. These findings underscore that the composite gradient, as a fundamental structural attribute, can significantly shape the functional interplay between different brain areas. This phenomenon was consistent with that observed in the macaque neocortex model [1], yet the mechanism remains to be elucidated.

**Fig. 6.**
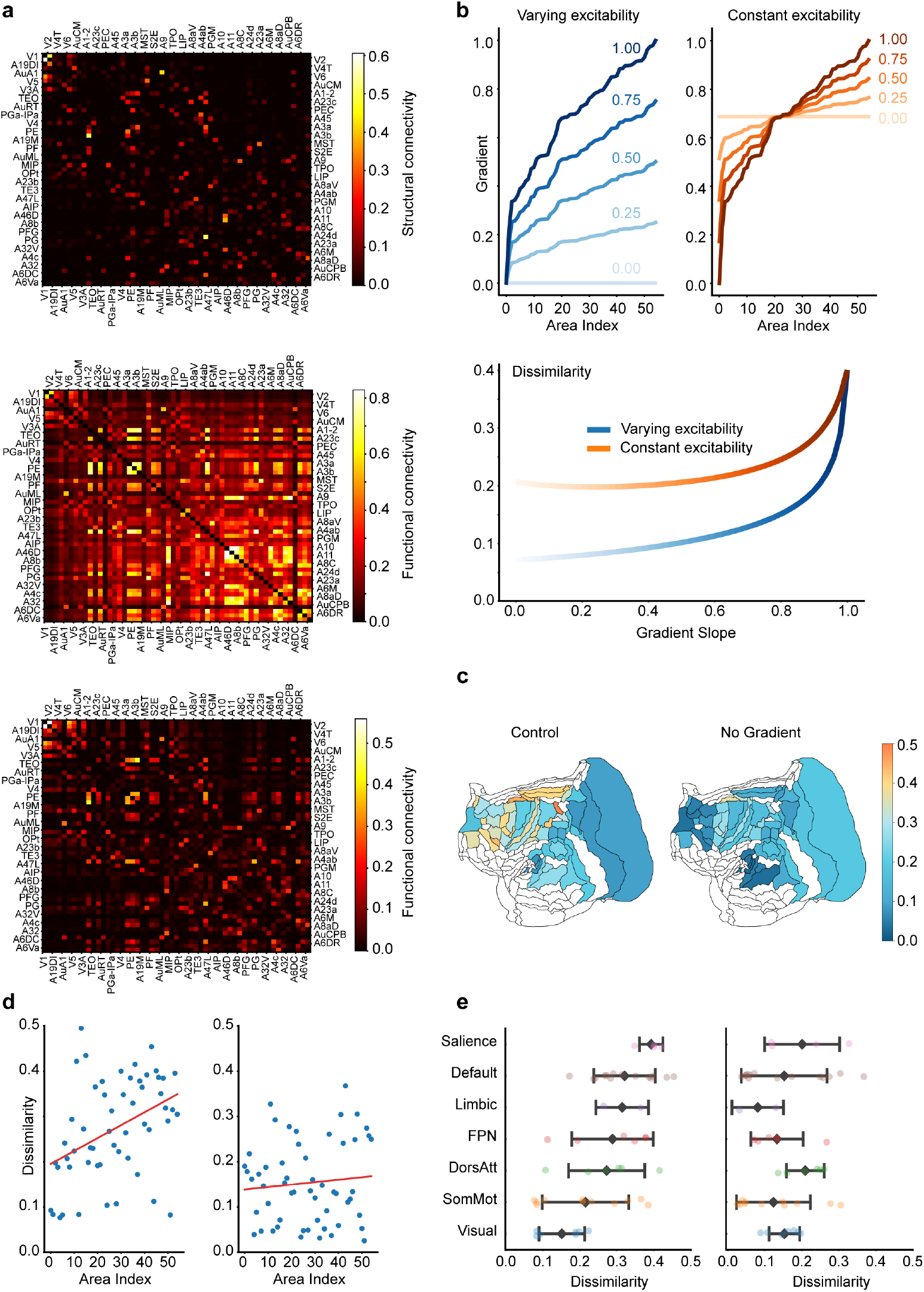
Impact of critical state on structural and functional connectivities. (a) Depiction of structural connectivity (SC) and functional connectivity (FC) under various scenarios. Top panel: Plot representing structural connectivity (SC) as per the FLN matrix. Middle Panel: Visualization of FC, measured by the correlation coefficient between resting-state neural activity from different areas, with the presence of composite gradient. Bottom Pane1: Visualization of FC in the absence of composite gradient. (b) Dissimilarity between FC and SC as a function of gradient slope. Top panel: Illustration of two distinct approaches to modifying the gradient slope, depicted through a color gradient from dark to light, indicating a change in slope from 1 to 0. Left, the adjustment of gradient slope with varying excitability. Right, the adjustment of gradient slope with constant average excitability. Bottom panel: The dissimilarity increases with the slope of gradient, with or without average excitability held constant. Here the dissimilarity is quantified as 1− |*ρ*| , where *ρ* is the Pearson correlation between FC and SC. (c) Heatmap of FC-SC dissimilarity across different brain areas, shown in marmoset brain parcellation. Left: FC-SC dissimilarity produced by the control model with composite gradient. Right: the model result with composite gradient removed. (d) Left, correlation between the FC-SC dissimilarity for each brain area ranked by its composite gradient of excitation in an ascending order (*r* = 0.41, *p* = 2.12 *×*10^−3^). Right, Absence of this correlation when the gradient is removed (*r* = 0.09, *p* = 0.491). (e) FC-SC dissimilarity across seven functional subnetworks of the marmoset brain. Left: the control model result. Right: The model result with gradient removed.

Here we further developed a mathematical analysis to identify a quantitative relationship between FC and SC and investigated their relationship when the macroscopic gradient of excitation exists, following the analysis in [38]. As shown in Figure 4a, the composite gradient gives rise to a wide range of timescale pool, in which the slowest timescale can be as long as hundreds of milliseconds or longer. The very slow timescale attributes to an eigenvalue close to zero, which indicates that the cortical network operates close to the critical state (the edge of stability). Building upon these facts, we mathematically analyzed the effects of criticality on the SC and FC relation (see Sec. Supporting Information), proving that the dissimilarity between FC and SC substantially increases when the network state approaches criticality.

We next scrutinized the role of the composite gradient on network criticality by adjusting its slope, akin to the model simulations conducted when investigating timescale localization. Specifically, we manipulated the slope of the gradients in two distinct ways: by either maintaining the average gradient value constant or allowing it to vary. Figure 6b (top panel) displays examples of these gradient adjustments for various gradient slope by each method. As depicted in Figure 6b (bottom panel), the dissimilarity between FC and SC intensifies with steeper gradient slope, confirming our hypothesis that composite gradients can contribute to the system’s criticality and, subsequently, the dissimilarity between FC and SC. We further explored the gradient’s function by maintaining the average excitability of all areas as a constant. Remarkably, even when the average excitability remains constant at the whole-cortex level, composite gradients can enhance local excitability and increase the dissimilarity between FC and SC (Figure 6b, bottom panel).

Next, we turned to analyze the impact of the composite gradient on individual cortical areas. Through mathematical analysis, we proved that the correlation between FC and SC is significantly lower in association areas compared to early sensory areas (see Sec. Supporting Information), which has also been reported experimentally in a monkey study [39]. In the model simulation, we computed the dissimilarity between FC and SC for each cortical area and visualized these differences in a marmoset brain parcellation (Figure 6c, left panel). The simulation results confirmed that dissimilarity is substantially higher in association areas than in early sensory areas. This trend is not evident when the composite gradient is absent (Figure 6c, right panel).

To further quantify this observation, we identified a positive and significant correlation between FC-SC dissimilarity and the rank of each area’s composite gradient (Figure 6d, left panel). In contrast, this correlation disappears when the gradient slope is zero (Figure 6d, right panel), indicating that this effect is not solely due to the heterogeneity of inter-areal connectivity. We further validated this by categorizing marmoset brain areas into seven subnetworks, comparable to those reported in humans [40] and macaques [41]. As shown in Figure 6e, FC-SC dissimilarity is relatively small for lower level visual and somatic motor subnetworks, but becomes more significant in higher level association subnetworks. These findings are consistent with previous findings on human cortex [41, 42] and are absent when the gradient slope is removed (Fig. 6e, right panel). This gradient-facilitated feature underscores the importance of higher-level areas in establishing wide-ranging functional connections, thereby supporting the execution of complex cognitive functions. These insights reaffirm the crucial role that gradients play in brain functionality.

The critical state of the model does not only depend on the composite gradient of excitation across areas but also on the inter-areal connection among areas. To study the effect of inter-areal connection on the FC-SC relation, we first removed the gradient from our model and studied the impact of increasing the inter-areal coupling strength of excitatory inputs. As the coupling strength escalates, the network state gets closer to criticality, resulting in an increased deviation between FC and SC. This trend, as depicted in Figure S4, is also consistent with our mathematical analysis (see Sec. Supporting Information).

### 2.6 Signal propagation in the near-critical state

In this section, we first investigated how the signal propagates across areas within our model, drawing parallels with observations from a previous optogenetics experiment [36]. In the experiment [36], an input was given to the marmoset A4ab area via an LED optogenetic stimulation and ECoG data were recorded from 64 electrodes broadly distributed on the marmoset neocortex. We replicated this experimental stimulus protocol within our model and analyzed the peak responses, defined as the highest level of neural activity observed in each cortical area post-stimulus. As shown in Figure 7a and Figure S6, our model well aligns with the experimental measurements for the majority of cases without additional parameter tuning, affirming the physiological relevance of our model’s responses. Similarly, this strong agreement with experiments was conserved after incorporating biologically realistic axonal conduction delays into our model (Figure S8).

**Fig. 7.**
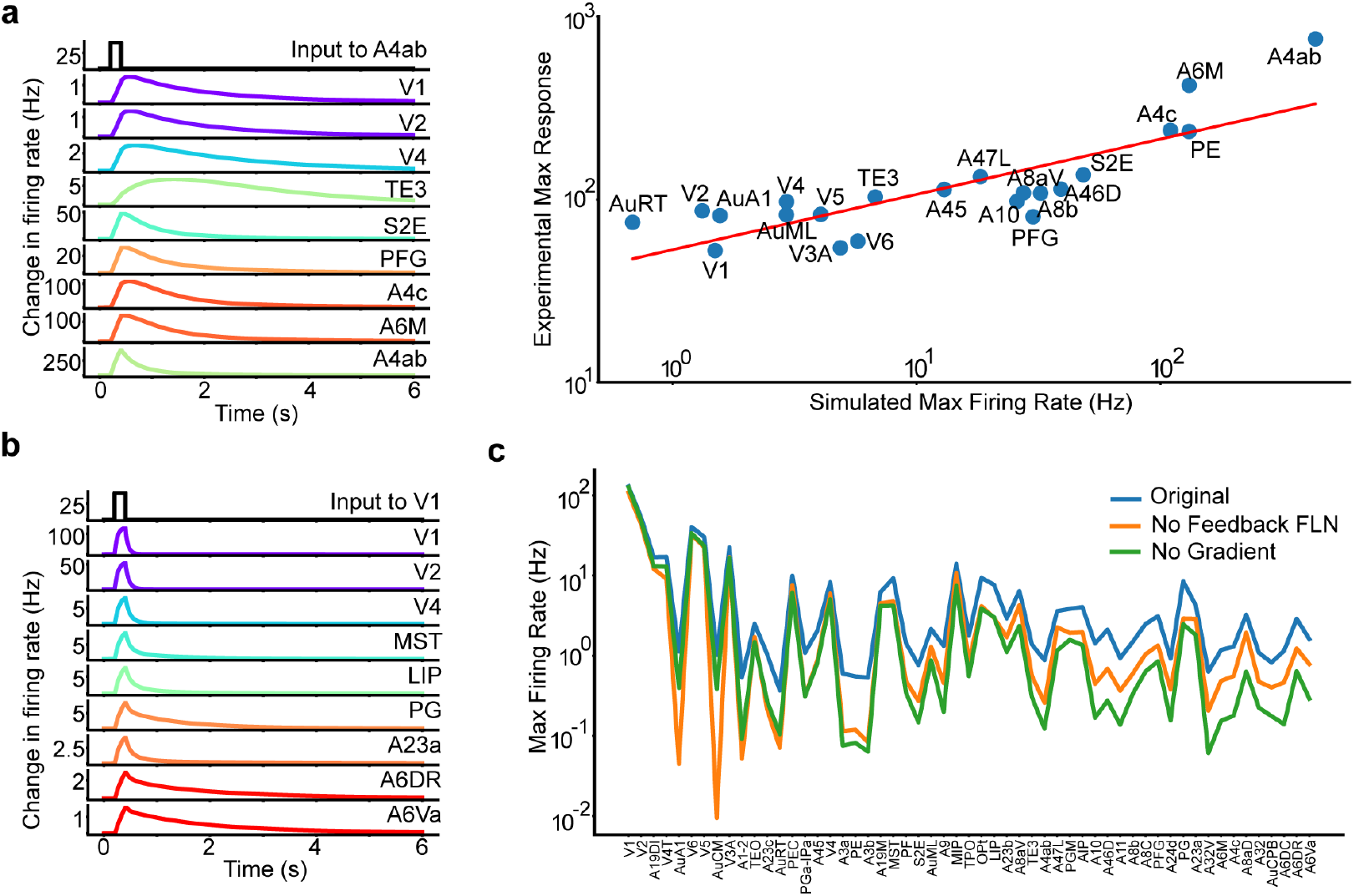
Signal propagation in the model. (a) Comparison of the model’s response post-stimulation to A4ab with corresponding optogenetics data. Left panel: Activity of representative areas following A4ab stimulation. Right panel: Comparison of the model’s peak response with optogenetics experimental data, demonstrating high model-experimental data consistency (*r* = 0.82, *p* = 3.21 *×*10^−6^). (b) Activity of representative areas following V1 stimulation. (c) Comparison of the model’s response in the control condition with the scenario when the feedback loops within FLN are eliminated and the scenarios where the composite gradients are removed.

We then examined the necessary conditions for signal propagation within the model starting from a pulse stimulus applied to sensory cortices, such as the primary visual cortex (V1), a central source of visual input. Responses from critical cortical areas were depicted in Figure 7b, showing steady signal propagation from the visual cortex to high-level areas like the prefrontal cortex.

We next investigated the role of inter-areal connectivity for signal propagation. The connectivity can be classified into feedforward and feedback connections based on anatomy [25, 26]. As intuitively expected, eliminating feedforward connections significantly hampers signal propagation from sensory areas (data not shown). We then shifted our focus to the role of feedback loops. These loops consist of connections from high-level areas to low-level areas identified by a smaller fraction of neurons in a projection originating from the supragranular layers of the source area (SLN *<* 0.5) [25, 26]. As demonstrated in Figure 7c (orange line), the deletion of all feedback loops within the FLN precipitates a generalized decrease in peak responses across all areas, with low-level areas bearing the brunt of this impact. Subsequent analyses investigated the influence of the composite gradient of excitation on signal propagation. As shown in Figure 7c (green line), the absence of the composite gradient led to a significant reduction in neural responses, particularly in higher-order areas. To explore this effect further, we systematically varied the gradient slope and examined the resulting changes in signal propagation, using peak responses from the PG area as a metric. As illustrated in Figure 8a, increasing the gradient slope enhanced signal strength, highlighting the critical role of gradients in facilitating signal propagation.

**Fig. 8.**
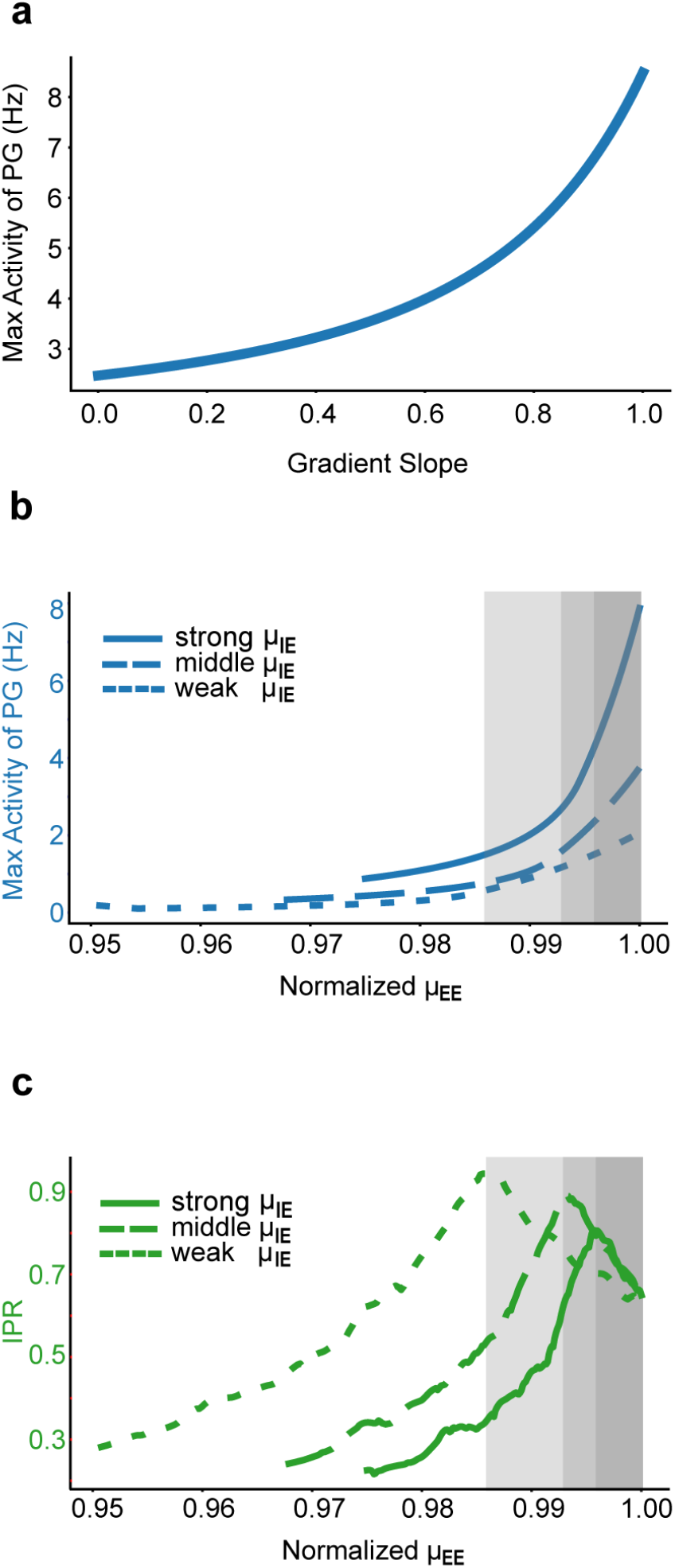
Signal Propagation and the critical state. (a) Illustration of a typical area (PG)’s response as a function of the slope of the composite gradient of excitability. (b)-(c) Impact of global excitatory connectivity (*µ*_*EE*_) on signal propagation and integration, where *µ*_*EE*_ is normalized to its critical value – the maximum at which the system retains stability. Linetypes indicate different *µ*_*IE*_ values: 49.81, 37.36, 24.91 pA/Hz for strong (solid line), middle (dash line), weak (dot line) *µ*_*IE*_ , respectively. Gray shaded area: the parameter region where exists the trade-off between signal propagation and integration. (b) Effect of *µ*_*EE*_ on signal propagation, quantified by the maximum activity in area PG’s response. (c) Effect of *µ*_*EE*_ on signal integration, quantified by IPR.

The contrast in signal propagation effects between the removal of excitation gradient and feedback connections is notable. The composite gradient of the excitation enables the high-level brain areas to receive stronger recurrent excitatory input, substantially amplifying long-range inputs from lower-level areas. In contrast, feedback connections enable indirect interactions between different lower-level areas via topdown projections from higher-level areas, significantly benefiting signal propagation, especially between areas lacking direct connections.

Building on the central role of both feedback loops and composite gradients in driving the system toward a critical state, our study delved deeper into the interplay between system criticality and the coexistence of signal propagation and integration. We examined the effects of altering global excitatory-excitatory (E-E) connectivity (*µ*_*EE*_), while maintaining a constant global excitatory-inhibitory (E-I) strength (*µ*_*IE*_), on the signal propagation and integration. Our findings, presented in Figure 8b-8c and Figure S7, elucidate several key insights. Firstly, the criticality of the system, implemented by the balance between *µ*_*EE*_ and *µ*_*IE*_, emerges as an important determinant of neuronal activities. An imbalance, particularly at lower *µ*_*EE*_ levels, markedly impairs both signal propagation and integration. Furthermore, we observed a tradeoff regime (grey areas in Figure 8b-8c, Figure S7) when *µ*_*EE*_ approaches the critical threshold, highlighting the intricate balance between signal propagation and integration. Approaching this threshold enhances signal propagation and enlarges the range of timescales, albeit at the expense of reduced timescale localization. Additionally, our results demonstrate that under conditions of strong global interaction strength (solid lines in Figure 8b-8c, Figure S7), both signal propagation and integration sustain high baseline values within the critical zone. This suggests that strong global connectivity, nudging the system toward criticality, is crucial in facilitating the co-existence of global signal propagation and localized integration.

## 3 Discussion

We reported empirical evidence for a hierarchy of time constants in the marmoset cerebral cortex and developed a connectome-based model of the marmoset neocortex, both for the first time to our knowledge. The key findings are threefold. First, a macroscopic gradient of synaptic excitation across areas, a detailed balance between excitatory and inhibitory populations, and the arrangement of long-range connections are instrumental in achieving timescale localization. Second, the state of the network operates close to criticality, an observable manifestation of which is the substantial dissimilarity between structural and functional connectivities, thus shaping the functional interaction between diverse areas of the neocortex. Third, our model accounts for experimental observations of signal propagation dynamics, underscoring the importance of the criticality facilitated by the feedback loops and the macroscopic gradients.

Criticality, or the edge of stability, is a state that can be signified by the system’s largest eigenvalue approaching zero. In the cortical network, it unveils itself as a powerful influencer of cortical functionality. This unique state naturally is closely linked to an enlargement of timescales, enabling signal integration with longer timescales than characteristic neuronal membrane timescales (hundreds of milliseconds versus tens of milliseconds). Moreover, criticality, achieved not solely through the balance of local excitatory and inhibitory inputs but also through broader network dynamics, facilitates extensive communication between local areas and the global neural network. This in turn amplifies signal propagation, affirming the vital role of criticality in streamlining neural communication.

Converging evidence across multiple species and brain regions indicates that cortical dynamics operate near a critical point, yet remain slightly subcritical [43–45]. This slight subcriticality is exemplified by a consistent branching ratio of ∼0.98 observed in vivo [45] and truncated neuronal avalanche size distributions in awake cortex [43, 44]. Notably, the precise operating point can be modulated by brain states: focused attention shifts cortical activity further into the subcritical state from the near-critical state during resting [46], while changes in arousal or anesthesia can also influence the distance from criticality [47, 48]. Operating just below criticality is hypothesized to confer several key functional advantages—ensuring stability against runaway excitation, preserving long-range correlations and large repertoires of activity patterns, enabling efficient information processing across a critical-like broad dynamic range, and allowing flexible tuning of network sensitivity [44, 49]. Theoretically, the cortex’s hierarchical modular architecture could yield an extended critical-like regime on the subcritical side (a Griffiths phase) that provides these functional benefits without finetuning [49]. Consistent with this “subcriticality” framework, our modeling results for the marmoset cortical demonstrate that a near-critical (slightly subcritical) state supports robust but controlled global signal propagation and naturally extended intrinsic timescales, highlighting how slight subcriticality effectively balances integration and stability while avoiding the fragility associated with exact criticality.

Our simulations and analysis further spotlighted the substantial impact of operating near criticality on the dissimilarity between Functional Connectivity (FC) and Structural Connectivity (SC). As criticality escalates, the disparity between FC and SC widens. This divergence permits the routing and broadcasting of information even in the absence of direct connections, promoting broad-range neural communication. Several pieces of evidence support the view that criticality contributes to this divergence in cerebral cortex. For instance, a prior model-based study showed that an enhanced criticality in the model could generate an FC more similar to experimentally measured FC [38].

Furthermore, recent studies on the human cortex have shown that the strength of structural–functional coupling (SFC) is not uniform across the cortex, but tending to be stronger in primary and unimodal sensory regions while progressively weakening in higher-order association areas [41, 42]. In line with these findings, our results demonstrate that applying a macroscopic gradient of excitation selectively modulates local criticality, enhancing signal integration and directing information flow in highlevel cortices. As a result, functional patterns in these higher-order areas deviate more strongly from their underlying structural connectivity compared to lower-level areas (Figure 6c), a divergence that may be crucial for supporting complex cognitive operations. Indeed, abolishing the gradient slope eliminates this spatially selective effect (Figure 6d), suggesting that it is not solely due to the heterogeneity of inter-areal connectivity. We further confirm that the decrease in SFC with increasing cognitive representational hierarchy remains evident when marmoset brain areas are categorized into seven subnetworks: SFC is strong in lower-level visual and somatic motor subnetworks but weakens in higher-level association subnetworks (Figure 6e). By tuning criticality through these macroscopic gradients, the brain may expand its range of potential activity states, thereby facilitating higher-order cognition.

Several elements of this study merit attention when compared with analogous previous modeling works on the macaque neocortex [1, 33]. Based on a similar modeling framework of cerebral cortex, which includes (i) representing each brain area by both excitatory and inhibitory group neurons; (ii) connectivity between areas dictated by experimentally measured connectomes; and (iii) a macroscopic area-wise composite gradient scaling the strength of excitatory projections, our model reproduces several phenomena observed in earlier studies—timescale localization, signal propagation, and the dissimilarity between FC and SC— despite modeling a different species, the marmoset. This conservation across species emphasizes the robustness of the model framework as a sturdy base for future research probing the relationship between structure and function at a whole-brain level, including the interplay between cortical dynamics and subcortical or peripheral neural systems.

Nevertheless, important distinctions from prior work are worth noting. Although both our study and earlier work constructed models with structures and parameters bound by experimental constraints, our study further corroborates the model’s responses with comparable experimental data across various scenarios, which allows us to compare the model and the experimental data area-by-area. We showed that the timescales for different areas predicted by the model and those measured in experiment under resting state (Figure 3c), and the after-stimulus response (Figure 7 and Figure S6) well agree with each other. The strong concordance with experimental data not only affirms the model’s validity but also accentuates its potential to probe intricate interactions within the cortical structural and functional dynamics.

Moreover, our model concurrently captures timescale localization (Chaudhuri et al. (2015)) and signal propagation (Joglekar et al. (2018)) with a single parameter set, which was not achieved in previous works. While Chaudhuri et al. (2015) revealed that the presence of a gradient can facilitate timescale localization, it significantly dampened signal propagation. Conversely, Joglekar et al. (2018) presented a model with satisfactory signal propagation but impaired timescale localization. Our model not only captures both phenomena within the same parameter regime but also exhibits similar qualitative behaviors to both models. This includes the gradient dependency for timescale localization, the necessity to maintain a balance between excitatory and inhibitory inputs, and the requirement of feedback loops for signal propagation. Understanding why our model could embody both features within a single parameter regime demands further mathematical analysis of this multi-regional model with gradients of excitation.

Our analysis also underscores a phenomenon wherein variations in model parameters yield diverse effects on timescale localization, signal propagation, and system criticality. For instance, increasing the gradient slope concurrently enhances timescale localization and signal propagation, nudges the system towards criticality, and amplifies the dissimilarity between FC and SC. Conversely, increasing the global coupling strength of excitatory neurons (*µ*_*EE*_) produces somewhat opposing effects. Although it boosts signal propagation and steers the system closer to criticality, thus increasing the disparity between FC and SC (Figure S4), it also undermines timescale localization. This reduction in timescale localization results from less isolated areas due to significantly enhanced long-range connections. Comparable impacts are observed when we adjust the strength of the FLN.

These diverse and somewhat paradoxical effects of various parameters highlight the complex interplay of factors that govern signal integration and propagation in the brain. Fully elucidating these intricate dynamics calls for more nuanced analyses and modeling work, going beyond the scope of our current study, to fully unravel the underlying mechanisms of brain functionality.

Another important consideration relates to the anatomical connectivity data used: the tracer-derived FLN weights currently employed in our large-scale marmoset cortical model do not specify neurotransmitter identity of target neurons [25, 26]. We therefore applied an identical FLN connectivity matrix to both excitatory (E) and inhibitory (I) neuronal populations within each cortical region, with their relative efficacies modulated by two global parameters, *µ*_*EE*_ and *µ*_*IE*_. These global scaling parameters were carefully tuned to simultaneously satisfy crucial dynamical constraints—notably, the observed hierarchy of intrinsic timescales and robust inter-areal signal propagation—and were subsequently kept fixed throughout all simulations. While this simplification enables effective modeling of largescale cortical dynamics, it neglects potential heterogeneity in cell-type specific projections. Future availability of cell-type resolved anatomical tracing data will refine the model by introducing projection-specific E/I targeting ratios instead of global scalar parameters.

While our model focuses exclusively on cortico-cortical connectivity, the cerebral cortex is not anatomically isolated – subcortical circuits (e.g., thalamus) and peripheral sensory pathways (retina, cochlea, spinal cord) impose important computational constraints on cortical activity. However, due to the lack of a marmoset-wide subcortical connectome at comparable resolution, our model necessarily focused on the corticocortical network, which can later be augmented with subcortical circuits as new data emerge. We anticipate that incorporating cortical-thalamo-cortical (CTC) and peripheral pathways in future marmoset models will refine our present findings, particularly by enhancing model predictions related to intrinsic timescale patterns, signal propagation, and functional connectivity profiles. Incorporating such pathways represents a key direction for future work as compatible anatomical datasets become available.

## Acknowledgments

We thank Cirong Liu for suggestions on area identification of the optogenetic ECoG public dataset, and thank Marcello Rosa and Sean FroudistWalsh for help categorize marmoset brain areas into seven subnetworks. This work is supported by National Key R&D Program of China 2023YFF1204200, National Natural Science Foundation of China Grant 12271361, 12250710674 and the Student Innovation Center at Shanghai Jiao Tong University (S.L.), James Simons Foundation Grant NC-GB-CULM-00003138 (X.-J.W.).

## 4 Methods

### 4.1 Experimental data sources

This work analyzed two experimental datasets to estimate neuronal activity’s time constants and signal propagation across cortical areas. The resting-state time constant was estimated from an ECoG dataset, gathered during an auditory task from a marmoset with a chronic implant in the left hemisphere [34]. The dataset features data from 96 uniformly placed electrodes throughout the hemisphere, gathered while the marmoset was in a resting state, under the influence of ketamine-based anesthesia (30 mg/kg i.m.). For the purpose of this study, we considered only the data collected prior to any auditory stimulus — approximately 15 seconds per trial — resulting in a cumulative total of 60 seconds of data compiled from four separate trials. We analyzed neural signals in 33 areas whose inter-areal connectivity has been measured in anatomical experiments [25, 26]. The activity of these areas can be directly modeled, which enables us to compare experimental data with simulation results.

For the analysis of signal propagation across areas, we utilized an optogenetic ECoG dataset collected from the right hemisphere of an awake marmoset [36]. This dataset was gathered using a combination of 64 epidurally implanted electrodes and eight LEDs designated for optogenetic stimulation. We utilized recording data with a single LED stimulating approximately areas A4ab, A6DR, A6Va, AuCM, S2E, V3A, PE, and PG. Each recording trial incorporated a 200ms pulse stimulus followed by a 2s post-stimulation interval, and the response was quantified by averaging the outcomes across a series of 50 trials.

For the intrinsic timescales during the awake resting state, we utilized the same optogenetic ECoG dataset for signal propagation analysis [36]. We specifically extracted the initial 5 seconds of recordings from the whole session — prior to any optogenetic stimulation—to obtain a resting-state segment free of task or laser effects. Nonetheless, due to the relatively short duration of awake-state recordings, we primarily used anesthetized-state data for subsequent analysis (e.g., comparisons with our model).

### 4.2 Estimation of the time constant of neural activity

We derived the time constants of neural activity during the resting state using a methodology based on Power Spectral Density (PSD) data [11]. This approach is especially beneficial when data length is short and the data contains oscillations and artifacts. The process entailed several stages. Initially, we computed the PSD using a variant of Welch’s method [50] that involved 1-second long Hamming windows with 0.5second overlap. For cases where multiple electrodes represented one area, we calculated the PSD individually for each electrode, subsequently taking the average as the PSD for that area.

Following this, we applied spectral parameterization [51] to decompose the PSD into a sum of Gaussians (representing temporal periodic terms) and a Lorentz function*L*(*f*) (representing temporal aperiodic terms) given by the form

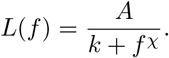

After fitting the data to determine the values of *A, k*, and *χ*, we identified the ‘knee’ frequency approximately as *f*_*k*_ = *k*^1/*χ*^. This frequency signifies the location where the PSD exhibits a ‘knee’ or bend. We then computed the corresponding timescale as 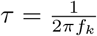.

For the post-stimulus activity timescale, we fitted the response following peak activity (post-stimulus) using an exponential function

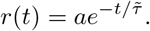

We determined the value of 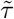 that provided the best fit as the time constant of the neural response in each cortical area.

### 4.3 Multi-regional cortical model architecture

We built the multi-regional marmoset cortical model using a set of ordinary differential equations, adapted from previous works of modeling macaque cortical network [1, 33]. In the model, 55 areas from sensory to association cortex were included. Each cortical area was modeled with one excitatory and one inhibitory neuron group governed by the following dynamics:

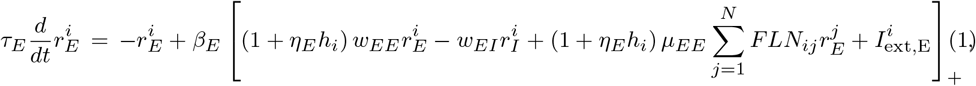

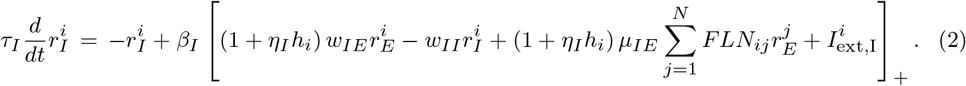

In these equations,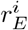 and 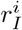 represent the firing rates of the excitatory and inhibitory populations in area *i*, respectively, while *τ*_*E*_ and *τ*_*I*_ are the corresponding intrinsic time constants. *w*_*XY*_ symbolizes the coupling strength from *Y* population to *X* population within the area (*X, Y* could be E or I population). *µ*_*XE*_ represents the coupling strength of the inter-areal input from the excitatory population to the *X* population in a downstream cortical area, while *FLN*_*ij*_ signifies the FLN from area *j* to area *i*. The external input to the *X* population in area *i* is expressed as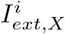 . We used a threshold linear function *f* (*x*) = *β*[*x*]_+_ for the gain function for both E and I populations, with gain factors *β* = *β*_*E*_ and *β*_*I*_ , respectively.

To complement our large-scale model that uses FLN to describe inter-areal connectivity, we additionally introduced a composite gradient of excitation, *h*. Because the FLN matrix is normalized per target area (each row sums to 1), it encodes only relative inter-areal connectivity but does not capture the area-wise differences in the absolute number of synapse inputs received by each target area. Anatomical evidence indicates that neurons in higher-order association cortex receive more inputs per cell than those in early sensory areas. For instance, dendritic spine counts increase systematically from sensory to association areas [26]. Furthermore, a uniform ∼80%*/*20% division of local versus long-range inputs across areas [52] implies that higher-order areas inherently possess greater numbers of both local and inter-areal synapses. Thus, neurons in association cortex receive a larger overall drive than those in lower areas. To account for this gradient, we define a composite gradient of excitation *h* as an area-specific scaling factor applied to all synaptic weights, effectively representing the increasing synaptic density in higher cortical areas while preserving the empirically measured FLN connectivity pattern. Within our model, *h*_*i*_ is a factor scaling both local (intra-areal) and long-range (inter-areal) excitatory inputs and varies across different areas. The quantitative estimation of *h*_*i*_ is detailed in the subsequent subsection. The scaling factor *η*_*X*_ governs the slope of the composite gradient on the *X* population.

Adapted from a previous experimental study [53], we set *τ*_*E*_ = 20 *ms, τ*_*I*_ = 10 *ms, β*_*E*_ = 0.066 *Hz/pA, β*_*I*_ = 0.351 *Hz/pA, η*_*E*_ = 0.685, *η*_*I*_ = 0.76, *w*_*EE*_ = 24.4 *pA/Hz, w*_*EI*_ = 19.7 *pA/Hz, w*_*IE*_ = 11.66 *pA/Hz, w*_*II*_ = 12.5 *pA/Hz, µ*_*EE*_ = 67.4 *pA/Hz, µ*_*IE*_ = 49.81 *pA/Hz*.

### 4.4 Incorporation of axonal conduction delays

We also extended the multi-area rate-based model to include explicit axonal conduction delays on all connections, both inter-areal and intra-areal for some studies. Inter-areal transmission delays, *τ*_*ij*_, were defined as proportional to the anatomical distance between the source area *j* and the target area *i*. Specifically, we set *τ*_*ij*_ = *d*_*ij*_*/v*_ax_, where *d*_*ij*_ is the white-matter tract length between areas *i* and *j* [25] and *v*_ax_ = 3.5*m/s* is the assumed axonal conduction velocity. All intra-areal (local) recurrent projections were assigned a fixed delay of 2*ms*. These parameter values align with those used in previous large-scale modeling studies of macaque cerebral cortex [33], thereby ensuring biologically plausible transmission latencies consistent with existing literature.

Delays were implemented by introducing a time shift in the firing-rate equations (Equations (1)-(2)) for each synaptic input. Specifically, any synaptic input from population *β* in area *j* to population *α* in area *i* was applied with a time offset corresponding to the assigned delay. For inter-areal connections (*i≠ j*), the presynaptic activity from population *β* at time *t*− *τ*_*ij*_ was used when computing input to the postsynaptic population *α* at time *t*. Similarly, for local recurrent connections (*i* = *j*), population activity at *t* −2*ms* was used in the coupling terms. This delayed-input mechanism was integrated the existing rate model dynamics without modifying any other parameters. The complete formula is as follows:

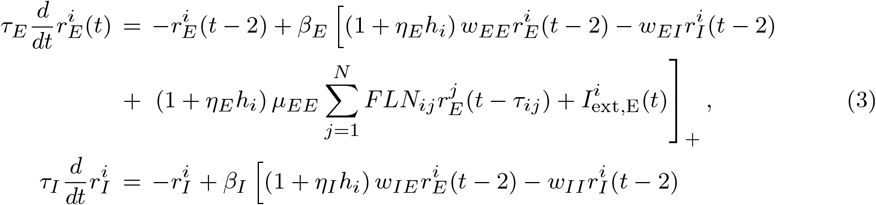

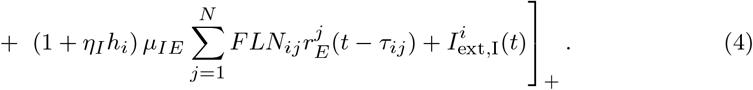

Importantly, introducing these biologically realistic conduction delays did not substantially change the primary results of the model: the global inter-areal sigal propagation patterns and the intrinsic timescale hierarchy remained robust and unchanged (Supplementary Figure S8).

### 4.5 Estimation of macroscopic gradient of excitation

We incorporated the composite gradient of excitation *h*_*i*_ in the model to be consistent with the experimental findings related to the gradient of timescale across areas and the structural hierarchy derived from the fraction of supragranular labeled neurons (SLN) as reported in Ref. [26]. As the model includes more areas (55 areas) than those recorded in the ECoG experiment (33 areas), the fitting was performed in two steps: firstly, we determined the composite gradient value of the 33 areas whose activity data and structural hierarchy data were both available. Secondly, we calculated the values of *h*_*i*_ for the rest areas based on the structural hierarchy and the values of *h*_*i*_ for the 33 areas obtained in the first step.

For the 33 areas whose timescale and structural hierarchy were both available, we used the following objective function that combined the similarity between the excitation composite gradient parameter *h*_*i*_ with the structural hierarchy *h*_*i,exp*_ determined by SLN and the timescale *τ*_*i,exp*_ derived from neural signals, i.e.,

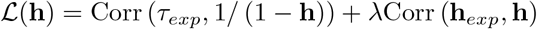

where **h**,**h**_*exp*_ and *τ*_*exp*_ are vectors, and *λ* is a hyper-parameter that determines the preference for the gradient to be closer to the timescale or the experimental hierarchy (chosen as 1 in this work). The Corr(*a, b*) is the Pearson’s correlation coefficient between *a* and *b*. We assessed the similarity with the structural hierarchy **h**_*exp*_ by measuring the correlation coefficient, and evaluated the similarity with the timescale *τ*_*exp*_ by calculating the correlation coefficient between 1*/* (1− **h**) and the experimentally measured timescale. We made this choice based on a mathematical analysis showing that 1 −**h** is proportional to the inverse of the time constant of neural activity when the multiple brain areas in the system are decoupled (details provided in Supporting Information). To obtain the values of the composite gradient, we solved the optimization problem presented by the objective function:

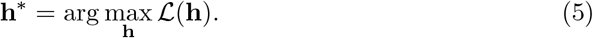

For areas whose timescale data could not be obtained from experiments (no electrode is placed in these areas), we determined their composite gradient *h*_*i*_ as a weighted average of the gradients of the 33 areas whose gradient *h*_*j*_ have been estimated in the first step. The weight was based on the similarity between structural hierarchy of twoareas:

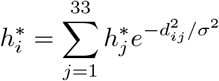

where *d*_*ij*_ =| *h*_*i,exp*_− *h*_*j,exp*_| is the difference of structural hierarchy for area *i, j* and *σ* = 0.05.

### 4.6 Simulation scenarios

We examined multiple simulation scenarios in the study. The scenarios encompassed a range of parameter configurations and manipulations of the network connectivity. For scenarios related to the change of gradient, we removed gradients (Figure 4a, 4d, 6a) by setting all *h*_*i*_ = 0. Furthermore, we manipulated the slope of the composite gradients by scaling with a factor *γ* (Figure 4b, 6b), leading to 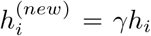 or by interpolating between the average gradient of all areas 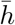 and the gradient *h*_*i*_ (Figure 6b)

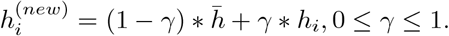

For Figure 6c, the gradient slope *γ* is set to 0.

In scenarios involving the change of network connectivity FLNs, we devised a condition devoid of any feedback loops by assigning zero to all FLN entries where the connection has a smaller fraction of neurons in a projection originating from the supragranular layers of the source area (Figure 7c), namely *FLN*_*ij*_ = 0, for all *SLN*_*ij*_ *<* 0.5. For Figure 5c to 5f, we shuffled the FLN by randomly permuting its entries 1000 times to obtain the distribution of the corresponding measures.

Regarding other parameter modifications, we disrupted the E-I balance by augmenting *w*_*EI*_ by 10% (Figure 5a). For Figure 8b to 8d, we set *µ*_*IE*_ = 24.91, 37.36, 49.81 *pA/Hz* , respectively and computed the corresponding upper bound 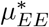 as 34.03, 50.68, 67.40 *pA/Hz* such that the system remains stable. *µ*_*EE*_ was varied as 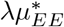 with *λ* ranging from 0.95, 0.97, 0.975 to 1 for each respective *µ*_*IE*_ setting. For Figure S4, we varied the value of *µ*_*EE*_ within the range from 64 to 70.

### 4.7 Metrics for timescale localization

We employed two quantitative measures to describe timescale localization across the whole neocortex. The first is the “inverse participation ratio” (IPR), which was originally introduced in quantum mechanics and solid-state physics. For a normalized eigenvector **v**, its IPR is defined as follows:

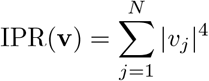

where *v*_*j*_ is the *j*th element of **v**. The IPR for the eigenvector matrix is computed as the average IPR across each column (eigenvector) in the matrix.

Furthermore, we introduced an additional metric for spatial localization of eigenvector **v** (Luis Carlos Garcia and Xiao-Jing Wang, see [37]):

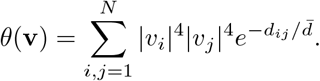

Here, *d*_*ij*_ represents the distance between a pair of areas *i* and *j*, which, in our context, denotes the physical distances between two areas in the marmoset neocortex; 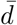 signifies the average distance across all pairs.

## 5 Data Availability

All data analyzed in this manuscript are publicly available from the Brain/MINDS project datasets [34, 36].

## 5.1 Code Availability

The source code and scripts used for simulation, analysis and figure generation are openly accessible on GitHub (https://github.com/GuanchunLi/MarmosetCortexModel/).

## Supporting Information

### Mathematical analysis of the relation between FC and SC

In this section, we presented an analysis on the relationship between functional connectivity (FC) and structural connectivity (SC) based on a simplified network model. We assumed the network only has excitatory neurons in each area with symmetric E-to-E connections. Despite this simplification, our analysis is able to conceptually explain the FC-SC relation observed in the original network model.

The dynamics of the *n*−dimensional system are governed by the equation:

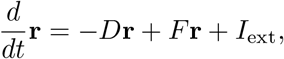

where *D* is a diagonal matrix representing the intrinsic timescale for each area and *F* is a symmetric inter-areal connectivity matrix. Both are assumed to be positive-definite matrices. The system can be re-written as:

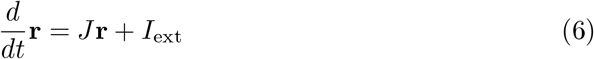

where *J* = −*D* + *F* and *J* = *J*^*T*^ . Notably, all eigenvalues of *J* must be negative for the system to remain stable.

In the resting state, when each area receives a white noise input of the same variance *σ*^2^ , we presented the following theorem for computing the functional connectivity (FC):

#### Theorem 1.

*The covariance matrix of* **r** *denoted by P satisfies the equation:*

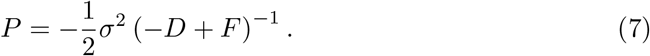

#### Proof.

Based on the analysis in Deco et al. (2013) [38], the covariance matrix *P* satisfies the Lyapunov equation:

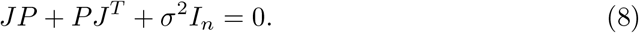

Since *J* is a real symmetric matrix, we can decompose it as

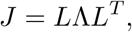

where *L* is a unitary matrix and Λ is a diagonal matrix with all diagonal elements being negative. By left-multiplying and right-multiplying (8) by *L*^*T*^ and *L*, respectively, we get

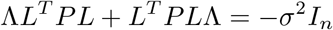

This leads to

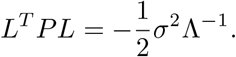

And thus,

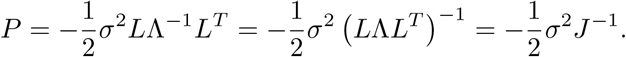

Substituting *J* = −*D* + *F* back into the equation, we have

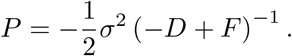

Based on the covariance matrix formula (7), we deduced the relationship between FC and SC as follows:

#### Theorem 2.

*When* ||*D*^−1^*F* || *<* 1,

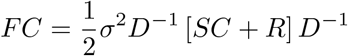

*with*

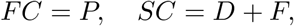

*and the remainder term is:*

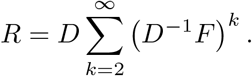

#### Proof.

From Theorem 1, we know that

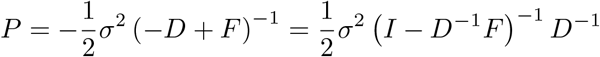

Given that 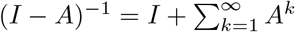,we have

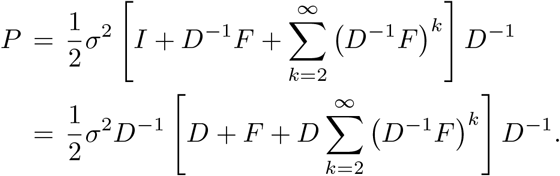

If we assign FC as *P* and SC as *D* + *F* (since it shares the same spatial structure with *J*), we find that

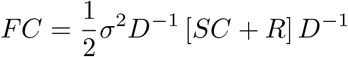

with the remainder term:

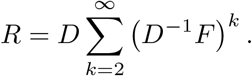

Theorem 2 suggests that the deviation between FC and SC can be fully encapsulated by the remainder term *R*. We then sought to understand how the value of *R* depends on the properties of the weight matrix *J*. We presented the following theorem:

#### Theorem 3.

||*R*|| → ∞ *as the system (*6*) approaches critical state, i.e*., *λ*_max_ (*J*) → 0.

#### Proof.

Considering the eigenvalue decomposition of *D*^−1^*F* as *D*^−1^*F* = *V* Ψ*V* ^−1^ with Ψ = diag (*ψ*_1_, *ψ*_2_, *· · ·* , *ψ*_*n*_), we know

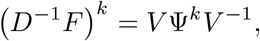

and

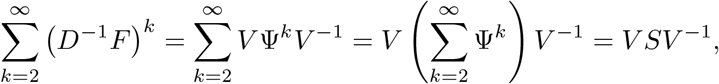

where

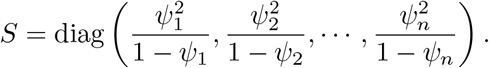

Hence we have

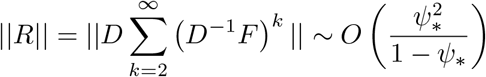

where *ψ*_*_ = *λ*_max_ is the maximum eigenvalue f *D*^−1^*F* . When *λ*_max_ (*J*) → 0, based on Lemma 1 below, we can have that *ψ*_*_ = *λ*_max_ *D*^−^ *F* → 1 and thus ||*R*|| →∞.

#### Lemma 1.

*As λ*_max_ (*J*) → 0, *it follows that λ*_max_

1. *Proof*. 1. First, we demonstrate that *λ*_max_ Given all eigenvalues of *J* = − *D* + *F* is negative, we know that *D* − *F* is a positive definite matrix. Based on the property that for two positive symmetric matrices *A, B* (Prob.III.6.14; Matrix Analysis, Bhatia 1997):

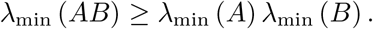

we have

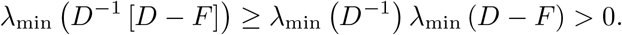 Since

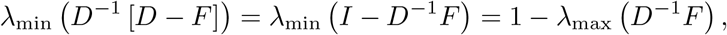

we have

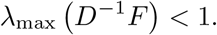
2. Next, we prove that *D*^−1^*F* contains an eigenvalue approaching 1 as *λ*_max_ (*J*) → 0. Letting *ϵ* = *λ*_max_ (*J*), there exists a corresponding eigenvector **x** such that

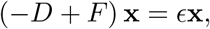

which is equivalent to

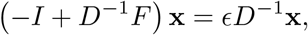

such that

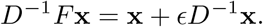

As *λ*_max_ (*J*) → 0, we know that *ϵ* → 0 and

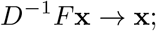

which indicates that *D*^−1^*F* has an eigenvalue approximating 1. Incorporating both findings that *D*^−1^*F* has an eigenvalue near 1 and *λ*_max_ *D*^−1^*F <* 1, we know that

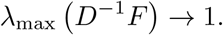

Furthermore, the corresponding eigenvector of *D*^−1^*F* also approximates to the eigenvector of *J*.

If the large entries (the magnitude being substantially nonzero) of the eigenvectors of *J* associated with near-zero eigenvalues are dense in higher indices (corresponding to areas of higher hierarchy), then the large entries of the remainder *R* are also densely concentrated within the block of higher indices. This can be shown by the following argument. Let us denote

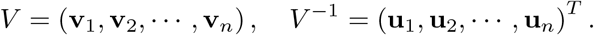

Based on the proof of Theorem 3, the remainder can be expressed as

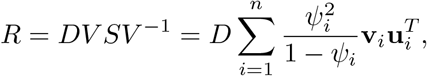

which gives out the relationship between the remainder and the corresponding eigenmodes of *D*^−1^*F* .

Our analysis and numerical results indicate that for the eigenvectors **u**_*i*_, **v**_*i*_ asso-ciated with eigenvalues *ψ*_*i*_ close to one, there will be prominent non-zero elements in **u**_*i*_, **v**_*i*_ corresponding to high-level areas. This observation stems from the understanding that the eigenmodes with larger time constants (linked to smaller eigenvalues for −*D* + *F* , which in turn correspond to eigenvalues closer to 1 for *D*^−1^*F*) often involve brain areas with a large gradient of excitation. Those significant non-zero elements in **u**_*i*_, **v**_*i*_ are further amplified by the factor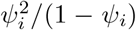 (as *ψ*_*i*_ is closer to 1), leading to large entries in the remainder *R*. This implies a considerable divergence between SC and FC in high-level brian areas. Such findings suggest that in systems exhibiting timescale localization, where higher-level areas possess larger timescales (corresponding to smaller eigenvalues), the discrepancy between SC and FC is more pronounced in these higher-level areas.

## Relationship between composite gradient of excitability and timescale of neuron activity

We derive the relationship between 1*/*(1 −*h*) (where *h* denotes the composite gradient of excitability) and the time constant of neural activity using a simplified version of the orginal model based on the Ornstein–Uhlenbeck process:

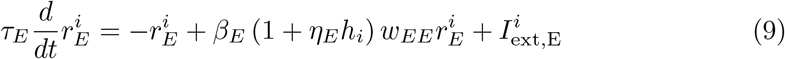

Here, the system, comprising both excitatory (E) and inhibitory (I) populations, is approximated by excitatory neurons alone, with the external input modeled as white noise 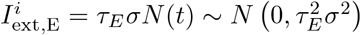.

The auto-correlation function of neural activity satisfying Eq.(9) can be solved as

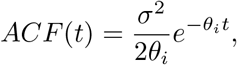

Where

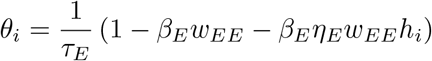

This yields the timescale of the neural activity as follows

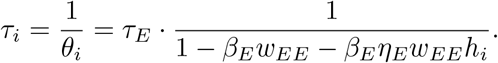

Thus, we have:

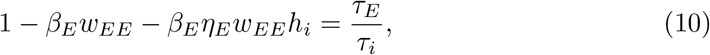

and consequently:

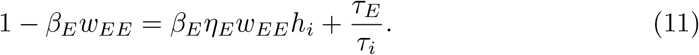

Notably, when the parameters are set in the model, for the high-level area with *h*_*i*_ → 1, the corresponding timescale *τ*_*i*_ is much larger (hundreds of milliseconds) than the intrinsic time constant *τ*_*E*_ (a few milliseconds), resulting in *τ*_*E*_*/τ*_*i*_ → 0. Incorporating this into Eq.(11), we have

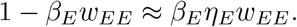

Taking this approximation back into Eq.(10), we obtain

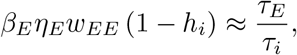

and hence

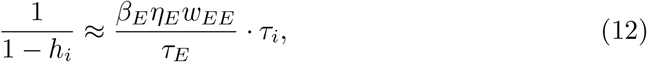

thereby establishing a linear relationship between 1*/*(1 − *h*) and *τ* .

### Stimulus-dependent timescale of neural activity

This section details our analysis of the network’s response following stimulus presentation to a specific brain area, particularly focusing on how the timescale of decaying activity depends on the stimulated area. Through mathematical analysis and numerical examples, we demonstrate that the observed discrepancies in timescales following different stimuli do not reflect a change in intrinsic timescale but rather the interactions among different brain areas.

We consider a linear system governed by the dynamics:

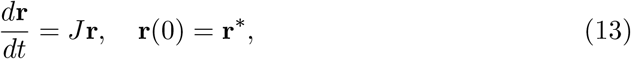

where **r**^*^ represents the system’s response post-stimulus, and we denote *t* = 0 as the time when the stimulus is off. Using the eigenvalue decomposition of the weight matrix *J*, we have

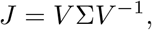

with eigenvalues *λ*_1_, *λ*_2_, *· · ·* , *λ*_*n*_ and corresponding eigenvectors **v**_1_, **v**_2_, *· · ·* **v**_*n*_ where

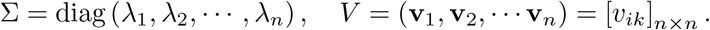

Projecting the firing rate vector **r** onto the eigenmode space yields

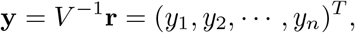

where each *y*_*i*_ denotes the projection magnitude onto the *i*-th eigenvector. The dynamics now becomes

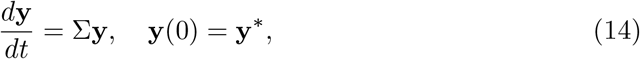

where **y**^*^ = *V* ^−1^**r**^*^, the initial firing rate profile projected onto the eigen-space.

Solving Eq.(14), we find

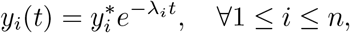

which leads to

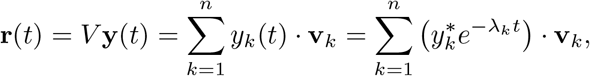

and thus

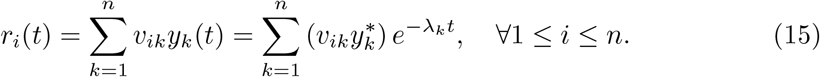

Eq.(15) illustrates that the dynamics of a given brain area’s activity are influenced by individual components with distinct timescales, determined by the term 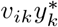,which combines the corresponding eigenvector element and the magnitude of **y**^*^ influenced by the response pattern of **r**^*^ post-stimulus.

Timescale localization ensures that *V* is a sparse matrix, with only a few nonzero elements for each *v*_*ik*_ per each given *i*. This sparsity indicates that each *r*_*i*_ will predominantly be influenced by a limited number of components, leading to distinct timescales across different brain areas. However, a component can still significantly influence the dynamics, even with a small *v*_*ik*_, provided that 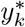 is substantially large.

This scenario is particularly likely when certain areas or clusters of areas are intensely activated during stimulus presentation.

Figure S1 illustrates the timescales and the contribution of each eigenmode to various brain areas following stimulus presentation, with either V1 or A4ab as the stimulated brain area. Following stimulation of V1, several eigenmodes associated with low-level visual areas are activated, typically characterized by shorter intrinsic timescales. In contrast, stimulation of A4ab predominantly activates eigenmodes involving a broader range of brain areas, including clusters of high-level areas, which generally exhibit much longer timescales. The activation of these eigenmodes becomes the dominant factor in the system’s behavior, significantly influencing the dynamics of low-level brain areas. This predominance explains why even low-level areas exhibit a slower decay in brain activity following stimulation.

In summary, stimulus presentation does not alter the intrinsic component patterns or the timescale of each component. Instead, it modifies the relative contributions of each component to different brain areas. This mechanism accounts for the observed discrepancies in timescales between the resting state and post-stimulus sceanrios. After stimulus presentation, a brain area’s timescale is influenced by inputs from other areas or clusters, particularly those strongly activated during the stimulus. This interplay results in the observed variation in decay time constants across different brain regions.

## 6 Supplementary Figures

**Fig. S1.**
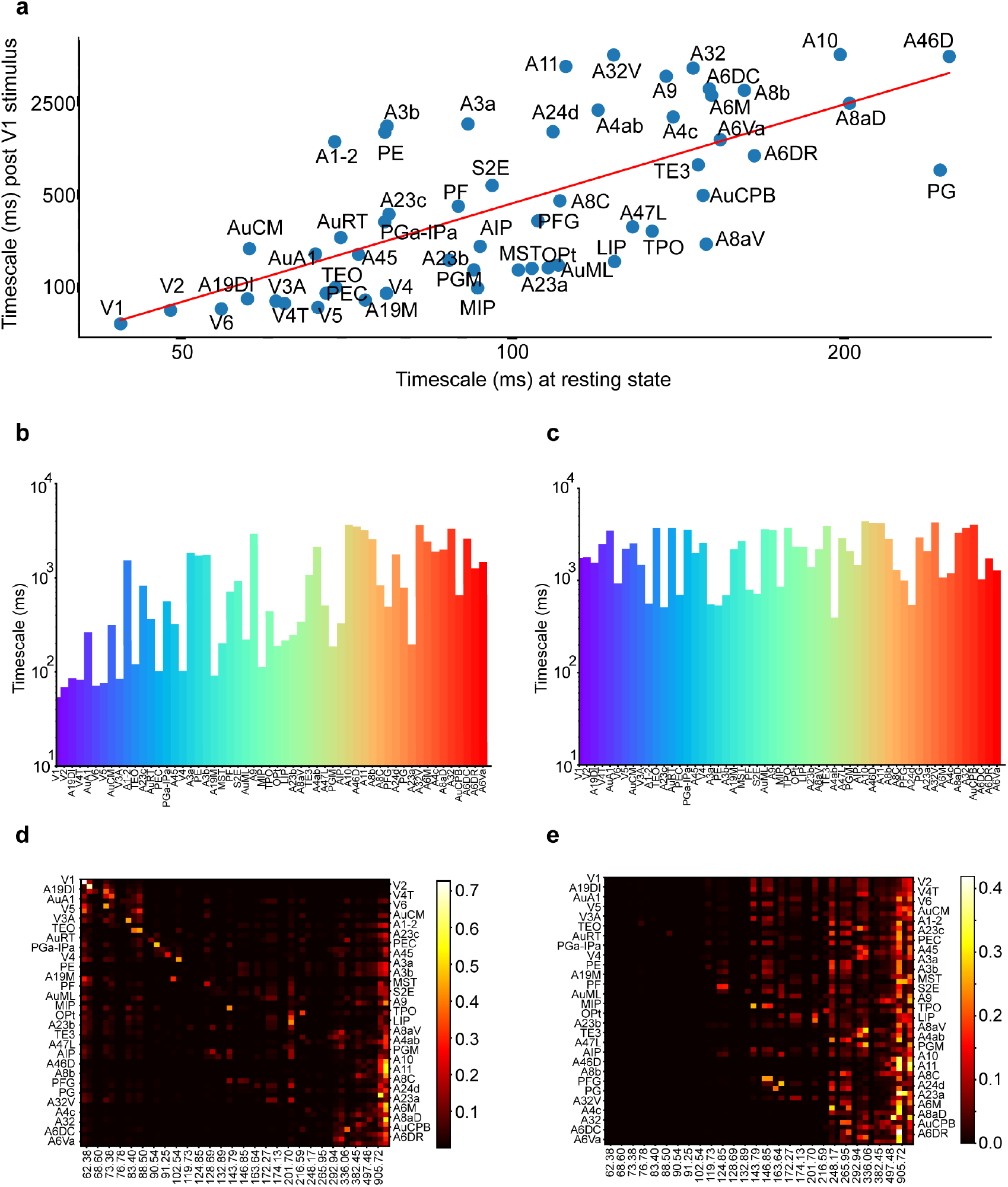
Stimulus-dependent timescale distribution across areas. (a) Comparison of timescale during the resting state and after V1 stimulus from model simulation. A strong correlation is observed (*r* = 0.69, *p* = 4.23*×* 10^−9^.) (b) The observed timescale gradient following a stimulus to V1, as extracted by fitting to the exponential function. (c) Similar to (b) but with stimulus to A4ab. (d) Relative contribution of each eigenmode to different brain areas (normalized by row) following stimulus presentation to V1. (e) Similar to (d) but with stimulus to A4ab.

**Fig. S2.**
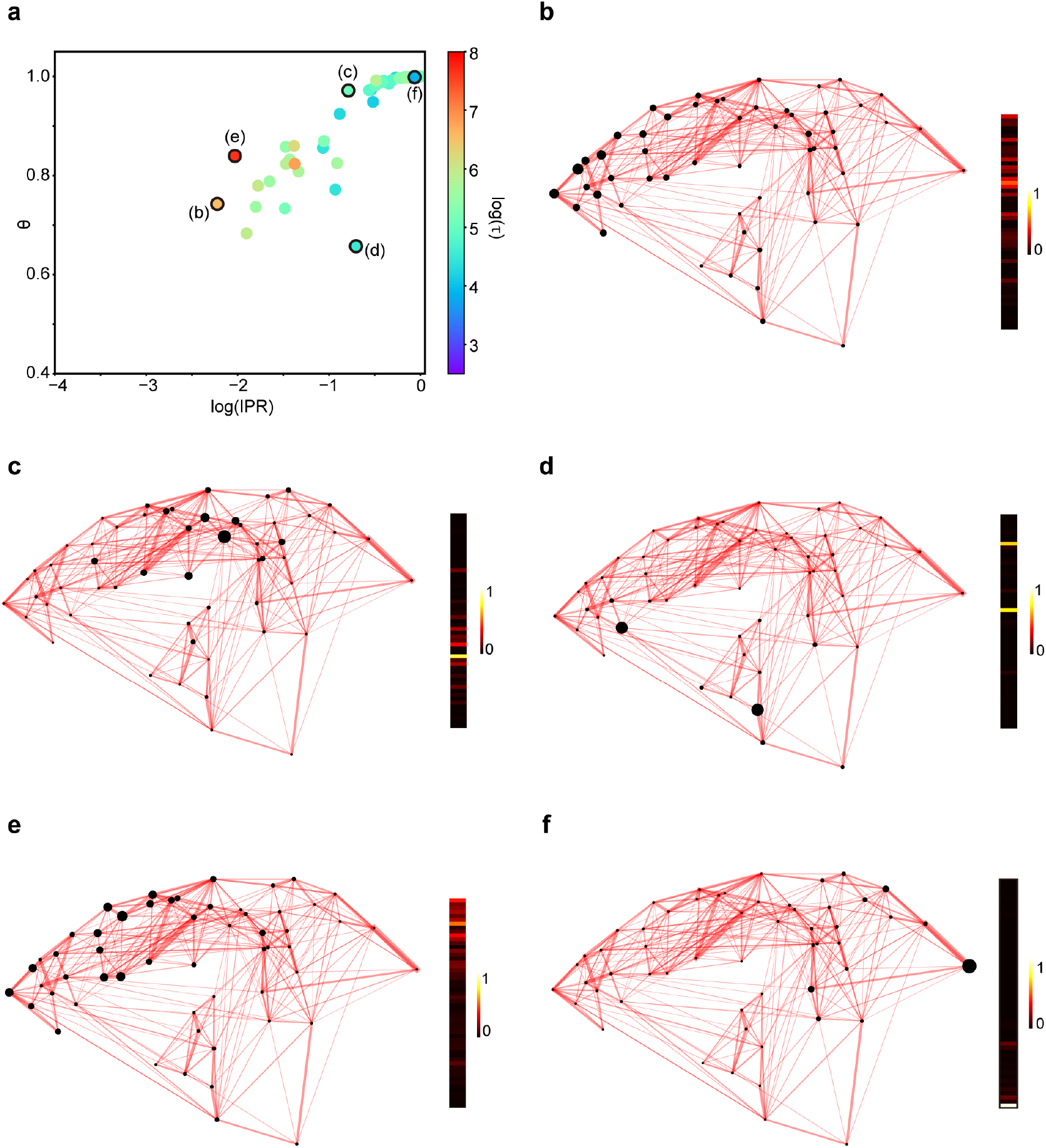
Two metrics to quantify timescale localization and spatial localization. (a) Scatterplot of IPR and *θ* for each eigenmodes of the model system. The color-coded timescale was after taken the logarithm using the natural base e. The five eigenmodes in (b)-(f) are marked with black edges. (b) - (f) Visualization of five representative eigenmodes illustrating timescale localization metrics IPR and *θ*, with values of eigenmodes plotted aside. The size of each black dot codes the magnitude of the corresponding element in the eigenmode. (b) An eigenmode with both weak timescale localization (IPR = 0.11) and weak spatial localization (*θ* = 0.74). (c) An eigenmode with both strong timescale localization (IPR = 0.45) and strong spatial localization (*θ* = 0.97). (d) An eigenmode with strong timescale localization (IPR = 0.49) but relatively weak spatial localization (*θ* = 0.66). (e) The slowest eigenmode, showing relatively weak timescale localization (IPR = 0.13) but relatively strong spatial localization (*θ* = 0.84). (f) Same as Figure 4c, the fastest eigenmode, showing both strong timescale localization (IPR = 0.91) and strong spatial localization (*θ* = 1.00).

**Fig. S3.**
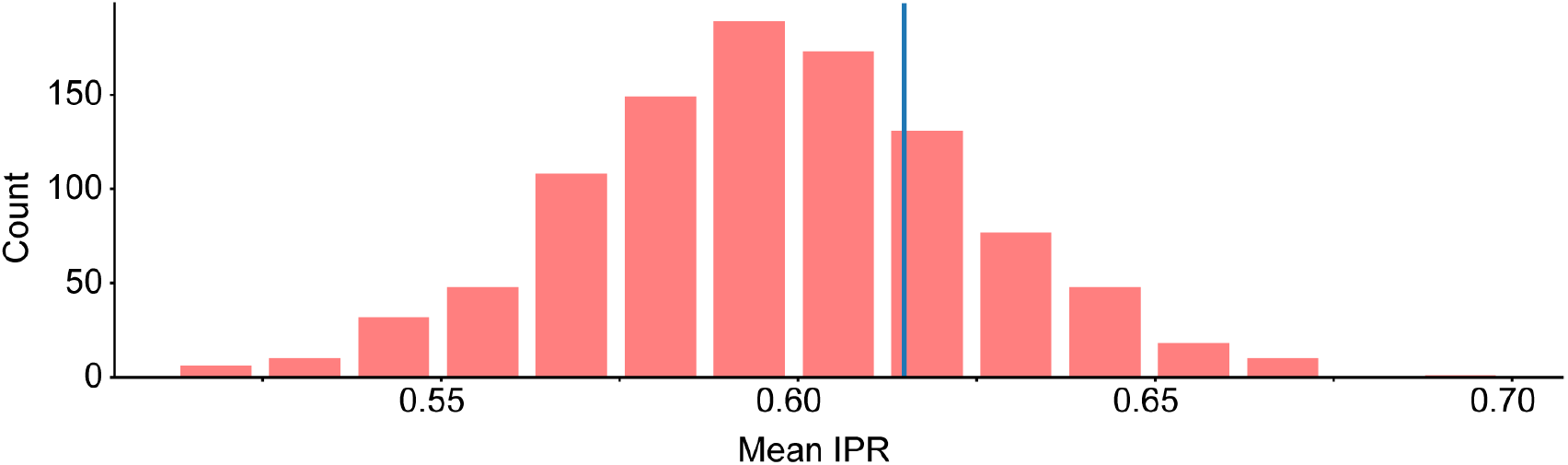
Impact of shuffled FLNs on timescale localization. The histogram of the degree of timescale localization quantified by the IPR metric with randomly shuffled FLN. The distribution is close to the value observed in the control condition, as indicated by the vertical line.

**Fig. S4.**
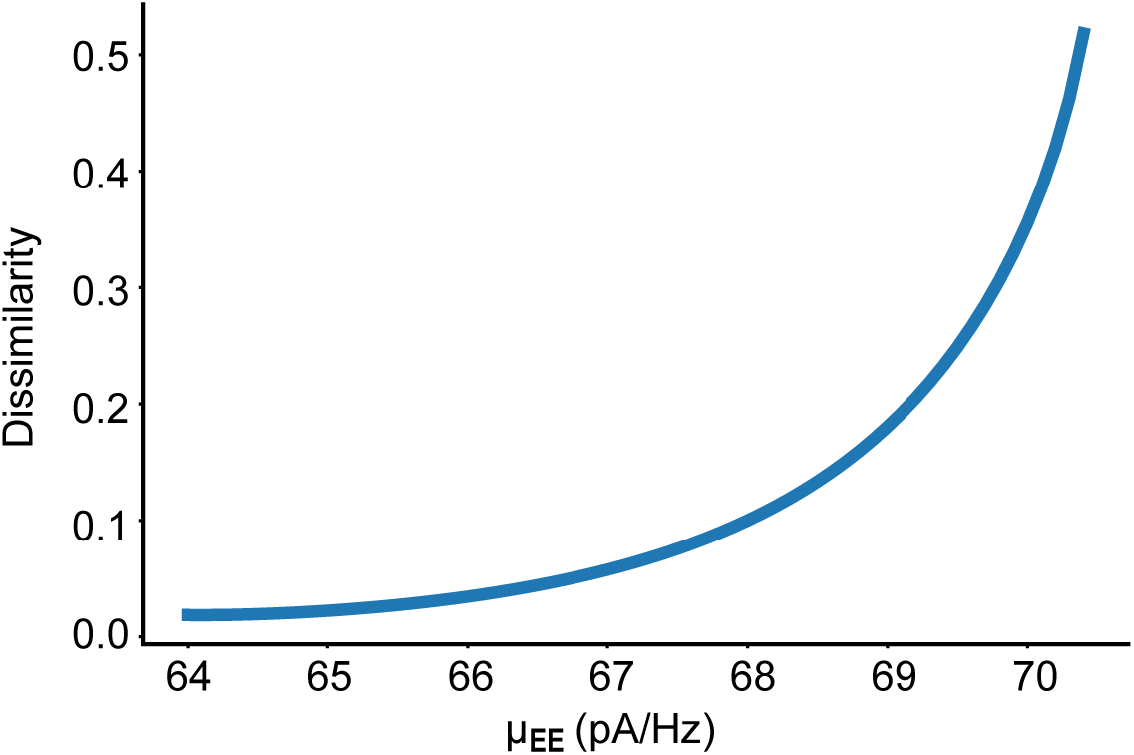
Dissimilarity between FC and SC as a function of global coupling strength *µ*_*EE*_ of excitatory neurons, another key parameter for criticality. Dissimilarity displays a monotonic increase with the strength of global coupling.

**Fig. S5.**
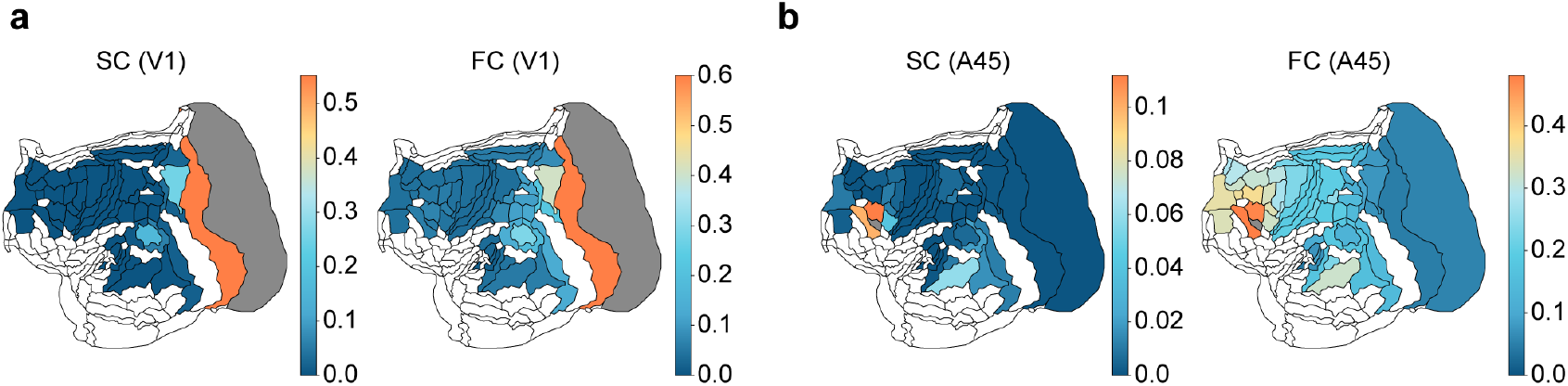
Comparison of structural connectivity (SC) and functional connectivity (FC) linked to different brain areas in marmoset brain parcellation. Left: Heatmap showing SC (left panel) and FC (right panel) of different brain areas connected to V1. Right: Heatmap showing SC (left panel) and FC (right panel) of different brain areas connected to A45.

**Fig. S6.**
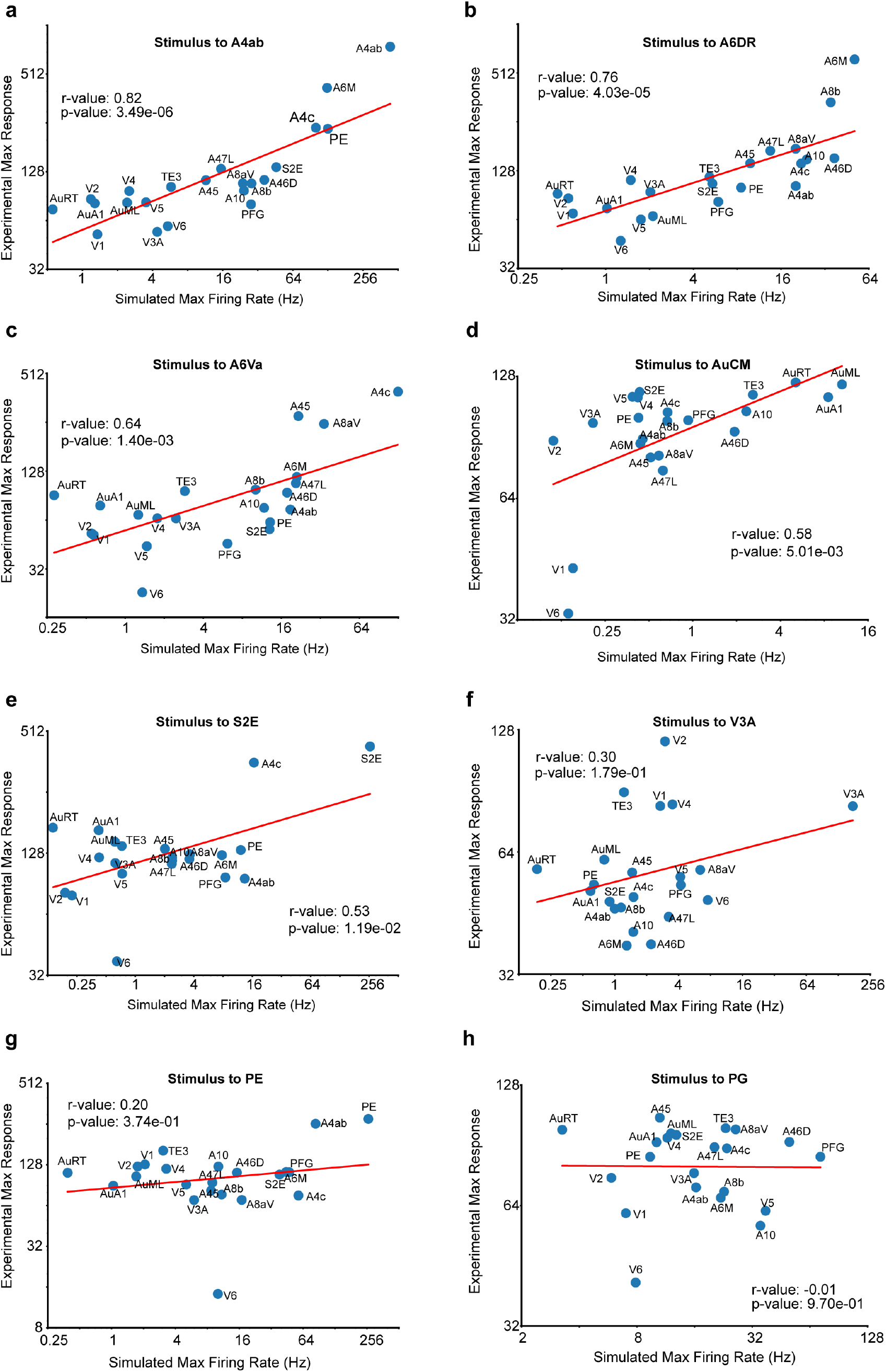
Comparison of the model’s peak response with optogenetics experimental data when different brain areas are stimulated, similar to Figure 7a. (a) - (h) Peak responses of multiple areas after stimulating putative brain area A4ab, A6DR, A6Va, AuCM, S2E, V3A, PE, PG, respectively.

**Fig. S7.**
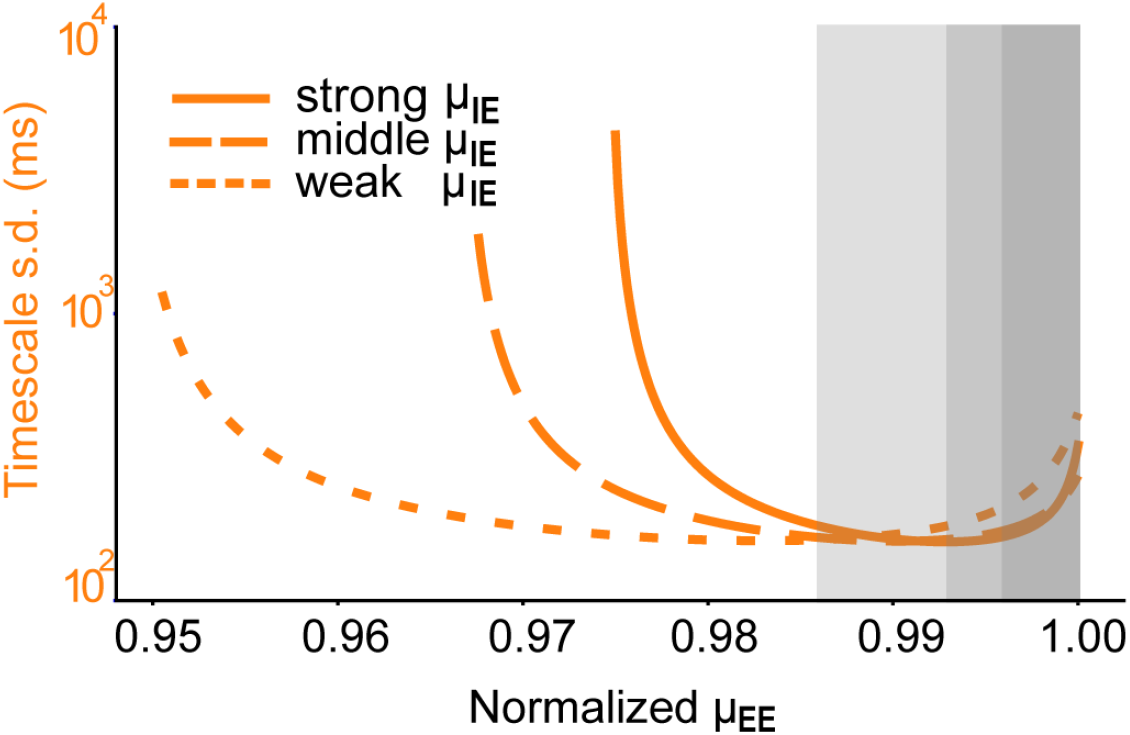
Impact of global excitatory connectivity (*µ*_*EE*_) on the timescale range, quantified by the standard deviation of timescales of different eigenmodes. The *µ*_*EE*_ is normalized to its critical value – the maximum at which the system retains stability.

**Fig. S8.**
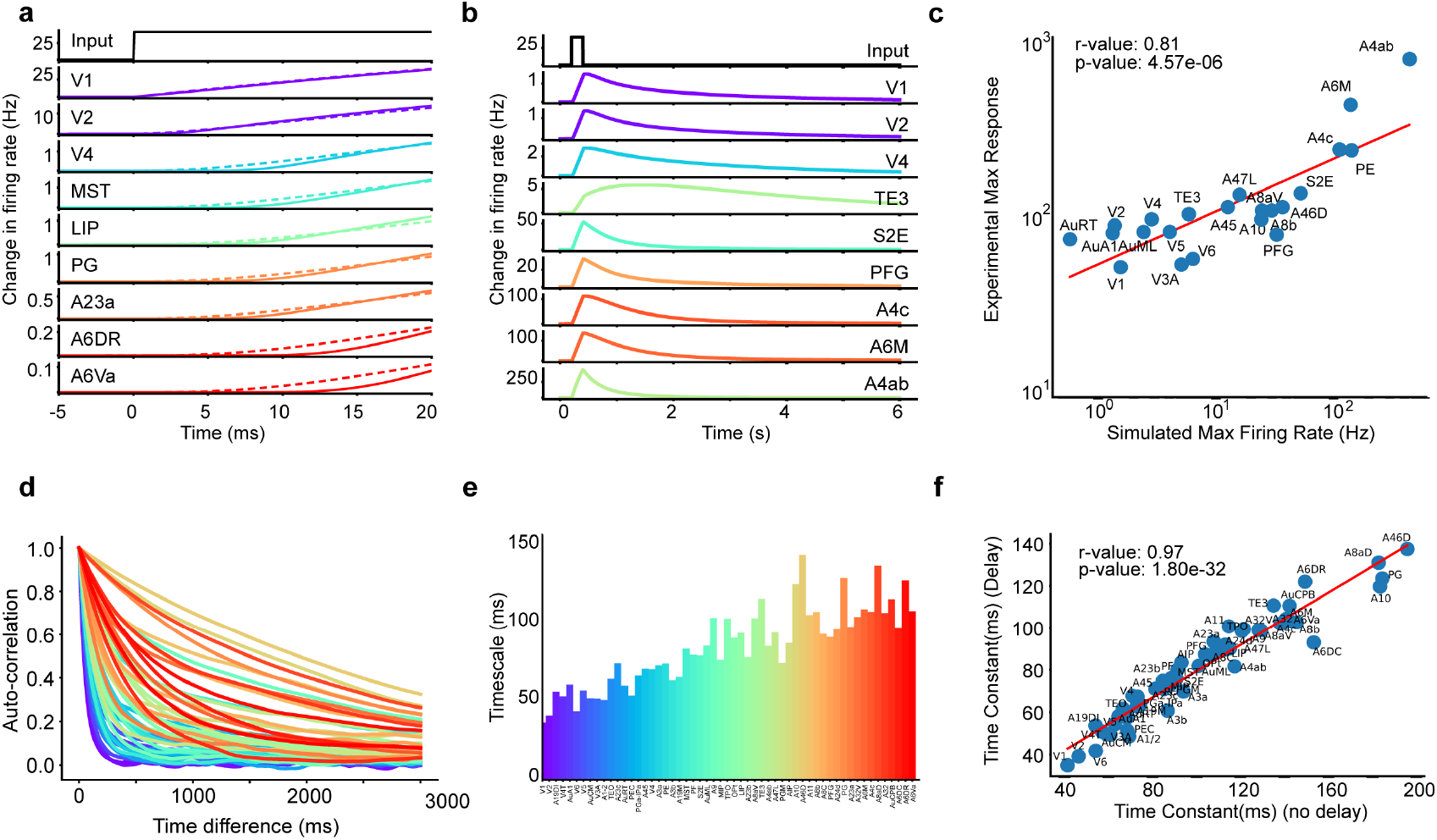
Results from the *model incorporating explicit axonal conduction delays*. (a) Activity of representative areas following V1 stimulation – focusing the first 20 milliseconds after stimulus onset. Solid line: the activity of model incorporating delays; Dash line: the activity of model without delay. (b) Activity of representative areas within the model incorporating delays following A4ab stimulation. (c) Comparison of the delay-incorporated model’s peak response with optogenetics experimental data, demonstrating high model-experimental data consistency (*r* = 0.81, *p* = 4.57*×* 10^−6^). (d) The autocorrelation function of each area’s activity in the resting state. (e) Hierarchical timescales extracted from simulated resting state activity of the model incorporating delays. (f) Direct comparison of intrinsic timescales between the delay-incorporated and delay-free models, demonstrating a robust consistence (*r* = 0.97, *p* = 1.80*×* 10^−32^) and indicating minimal impact of conduction delays on the overall timescale hierarchy.

**Fig. S9.**
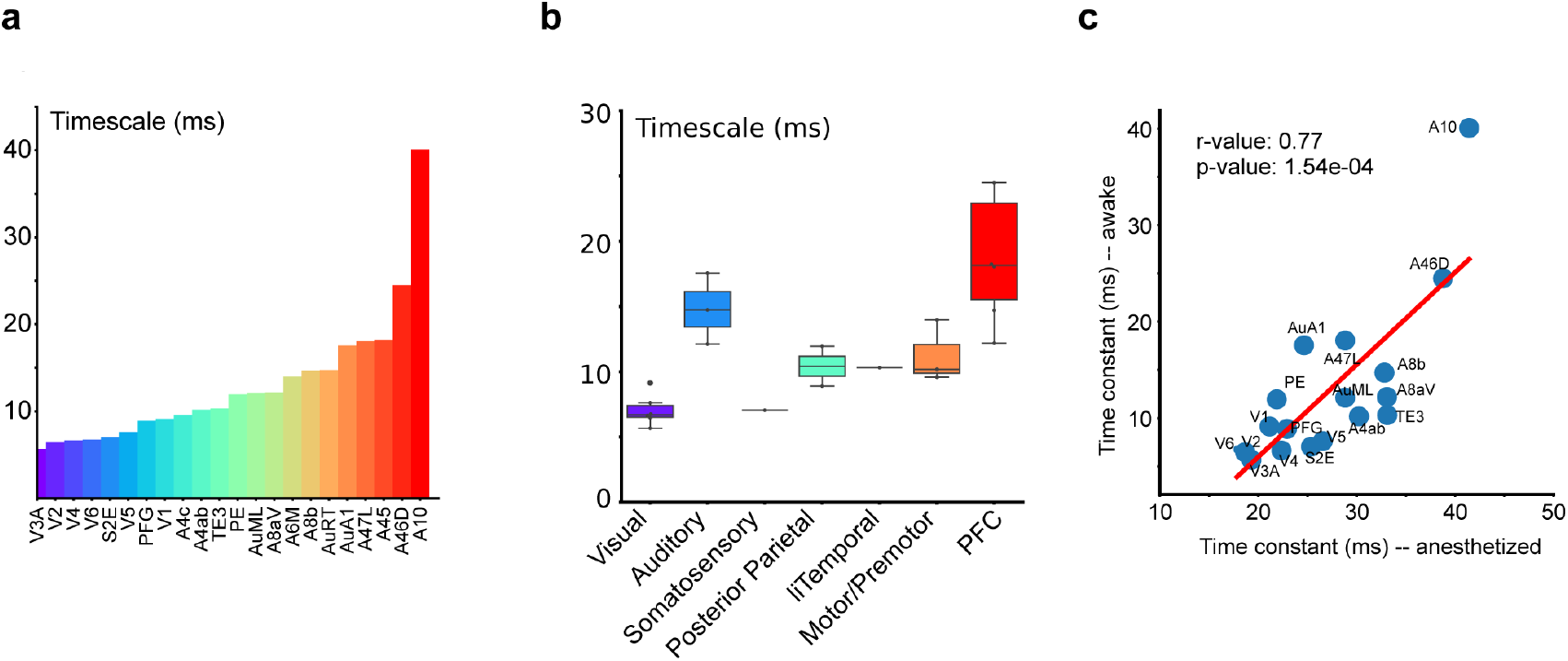
The timescale hierarchy in the marmoset cortex during the awake resting state. (a) Timescale distribution across the marmoset cortex, derived from neural activity recorded during the awake resting state. Lower-level sensory areas such as V1 and V2 exhibited shorter timescales, in contrast to higher-level areas like A46D and A10 that exhibited longer timescales. (b) Boxplot of timescale statistics for each areal category. Box plots show the media and IQR, with whiskers extending to the minimum and maximum values excluding outliers. (c) Strong correlation between timescales estimated from anesthetized and awake resting-state data (*r* = 0.77, *p* = 1.54*×* 10^−4^), confirming that the observed timescale hierarchy is consistent across brain states.

